# Metabolic adjustment of *Drosophila* hemocyte number and sessility by an adipokine

**DOI:** 10.1101/648626

**Authors:** Elodie Ramond, Bianca Petrignani, Jan Paul Dudzic, Jean-Philippe Boquete, Mickaël Poidevin, Shu Kondo, Bruno Lemaitre

## Abstract

In animals, growth is regulated by the complex interplay between paracrine and endocrine signals. When food is scarce, tissues compete for nutrients, leading to critical resource allocation and prioritization. Little is known about how the immune system maturation is coordinated with the growth of other tissues. Here, we describe a signaling mechanism that regulates the number of hemocytes (blood cells) according to the nutritional state of the *Drosophila* larva. Specifically, we found that the adipokine NimB5 is produced in the fat body upon nutrient scarcity downstream of metabolic sensors. NimB5 is then secreted and bind to hemocytes to down-regulate their proliferation and adhesion. Blocking this signaling loop results in conditional lethality when larvae are raised on a poor diet, due to excessive hemocyte numbers and insufficient energy storage. Similar regulatory mechanisms shaping the immune system in response to nutrient availability are likely to be widespread in animals.

**Author summary:** *Drosophila* larval hemocytes (blood cells) are found in two compartments: the lymph gland considered as a reservoir, and the peripheral compartment. Peripheral hemocytes form sessile patches attached to the internal surface of the larval body wall or are found freely circulating in the hemolymph. Little is known about the signals that regulate hemocytes proliferation and localization in the peripheral compartment. In this study, we have identified a new gene, *NimrodB5*, coding for the NimB5 protein, which is secreted by the fat body and binds to hemocytes. NimB5 inhibits hemocyte proliferation while promoting sessility, leading to an increased number of circulating hemocytes and adhesion defects in *NimB5* mutant. We show that *nimrodB5* expression by the fat body is controlled by metabolic cues to adjust hemocyte number to the physiological state of the larvae.

Interestingly, deregulation of *NimB5* causes lethality when larvae are raised on a poor diet due to a defect in regulating hemocytes proliferation. In conclusion, we have identified a new adipokine that optimizes hemocytes number to the physiological state of larvae. Our study also reveals a major role of the fat body in peripheral hematopoiesis regulation and outline how it can be costly to maintain a basal immune defense.

## Introduction

In multicellular organisms, growth is regulated by the complex interplay between paracrine and endocrine signals. Tissues and organs respond to specific genetic programs to ensure optimal animal development with harmonious proportions [1]. Environmental cues, notably nutrient availability, further modify the output of these programs. Under food scarcity, critical resource allocation and prioritization between the various tissues of the organism are required to sustain life. This condition can favor brain development at the expense of other organs or an optimal size, with long-term consequences on animal fitness [2].

The immune system forms a network of tissues and circulating cells whose primary function is to prevent and limit microbial infections. The development and maintenance of an immune system are metabolically costly. Theoretical considerations suggest that resource allocation towards the immune system is determined by the diversity of pathogens and the recurrence of infections, as well as by trade-offs with other physiological or reproductive functions [3–5]. Examples of trade-off have been shown in poultries where selection for increased body mass resulted in a decrease in immune function [6] or conversely in *Drosophila* where selection for increased resistance to parasitoid wasps resulted in reduced larval competitive ability on a poor diet [7]. Another illustration is the allocation of resources an animal will invest in the production of immune cells. In humans, extremely high numbers of circulating leukocytes are continuously generated to anticipate potential infections by pathogens. While low lymphocyte or neutrophil numbers have been associated with increased risk of infection, the excessively high blood cell counts in certain lymphomas cause exhaustion of host resources and sometimes death [8]. As opposed to many vital physiological functions, the immune system provides minimal benefits under basal condition, revealing its importance mainly in case of an infection. When nutrients are scarce, it is expected that resources are re-allocated to critical developmental functions at the expense of the immune system [9]. This re-allocation would explain why starved children are more vulnerable to infections [10,11]. It also suggests the existence of flexible mechanisms, which allocate resources to the establishment of an immune system depending on nutritional inputs. However, our knowledge about how immune system development is coordinated with that of other organs in case of nutrient scarcity is limited. In this article, we describe a mechanism that regulates the number of hemocytes (i.e. insect blood cells) according to the nutritional state of the developing *Drosophila* larva.

The *Drosophila* cellular immune response involves three types of hemocytes, two of them, plasmatocytes and crystal cells, are found in the absence of infection. Plasmatocytes represent the most abundant population (i.e. 90-95%) and share functional similarities with mammalian macrophages. Crystal cells are non-phagocytic cells, which synthesize enzymes and substrates responsible for producing melanin at an injury site or for fighting an infection [12,13]. When faced with infestation by parasitoid wasps, *Drosophila* larvae produce a third type of blood cells: the lamellocytes. These cell types differentiate from plasmatocytes [14] or hemocyte progenitors [15] and form a capsule around wasp eggs or intruders, which are too large to be phagocytosed.

Like in vertebrates, *Drosophila* hematopoiesis occurs in several waves. The first wave of hematopoiesis occurs during embryogenesis and gives rise to approximately 700 plasmatocytes and 30 crystal cells [16]. The embryonic hemocyte population expands until the third larval stage to stop at metamorphosis. Outside the hematopoietic organ, larval hemocytes are found both in circulation and in sessile patches (referred to as peripheral hemocytes) [17–23]. Sessile hemocytes are attached to the internal surface of the larval integument, forming patches, some of which are closely associated with secretory cells called oenocytes, as well as with the endings of peripheral neurons [19,24]. Larval hemocytes continuously swap between sessile patches and the hemolymph compartment [25,26]. The second wave of hematopoiesis takes place in a dedicated organ called the lymph gland [17,18]. The lymph gland functions as a reservoir of hemocyte progenitors and mature hemocytes, releasing hemocytes at the onset of metamorphosis or upon parasitic infestation. The hemocyte population in pupae and adults consists of a pool of both embryonic and lymph gland-derived plasmatocytes with rare crystal cells [27].

In this study, we investigate whether peripheral hematopoiesis is influenced by nutrient scarcity. We first demonstrate that excessive hemocyte numbers can be detrimental under conditions of nutrient deprivation since they prevent the deposition of lipids in the fat body, which is the primary energy storage organ. We then show that peripheral hematopoiesis is down-regulated under poor dietary conditions, suggesting the existence of a regulatory pathway that adjusts hemocyte number to the metabolic state of the host. Finally, we identified an adipokine, NimB5, which is secreted by the fat body upon nutrient deprivation and which promotes hemocytes sessility and reduces their proliferation. Blocking this regulatory loop prevents the adjustment of hemocyte numbers to the nutrient state of the host, resulting in lethality under restrictive diets.

## Results

### High hemocyte numbers are detrimental to larva upon nutrient scarcity

Production and maintenance of hemocytes are likely to have a significant metabolic cost. Therefore, we hypothesized that excessive hemocyte numbers might negatively impact *Drosophila* development. To test this hypothesis, we compared the viability of larvae with normal, higher or lower hemocyte numbers when raised on a rich or a poor diet. Larvae with increased hemocyte numbers were obtained by over-expressing *pvf2* (UAS-*pvf2*) [28] or *ras* (*UAS-ras85D^V12^*) specifically in plasmatocytes with the *hml^Δ^GAL4,UAS-GFP* driver, or using *eater^1^* larvae, deficient for the Eater transmembrane receptor [29,30]. Flow cytometry measurements on third instar (L3) wandering larvae showed that hemocyte numbers increased by 8.4- and 20.8-fold when over-expressing *pvf2* or *ras85D^V12^*, respectively, compared to wild-type when raised at 29°C. e*ater^1^* larvae exhibited about 3-fold higher hemocyte numbers than wild type larvae when raised at 25°C (Figures 1A, B). Larvae with reduced hemocyte counts were obtained by silencing *ras* in plasmatocytes, resulting in 3.6-fold lower hemocyte counts (Figure 1A). We noted that, independently of their hemocyte numbers, all larvae developed well when raised on a ‘rich’ standard medium, with the emergence of wild-type-like adults. Strikingly, all larvae with higher hemocyte numbers died when raised on a poor diet, with earlier lethality for mutants having the highest hemocyte numbers (> *pvf2* and *>ras^V12^* at L1/L2 and *eater^1^* at the pupal stage) (Figures 1C, D).

In contrast, wild-type or *ras RNAi* larvae with average or lower hemocyte numbers developed normally when raised on a poor diet. A way to explain our results is that larvae with higher hemocytes counts are in an “inflammatory-like” state that is deleterious to the host. Indeed, chronic activation of the immune system has been associated with precocious lethality in *Drosophila* [31,32]. Therefore, we monitored in these larvae the level of activation of the Imd, Toll and JAK-STAT immune pathways using appropriate read-out genes *(Diptericin, Drosomycin*, and *upd3* respectively). We did not observe any chronic activation of these pathways nor defect in immune inducibility in larvae with higher hemocyte count (Figure S1A-C). Moreover, hemocytes from larvae over-expressing *pvf2* or *ras^V12^* retain the ability to phagocyte Gram-negative and Gram-positive bacteria (Figure S1D, E). These data indicate that hemocytes from larvae with higher blood cell count are fully functional and the lethality observed on a poor diet cannot be explained by the chronic activation of the immune system. Previous studies have shown that over-proliferating tissues can induce energy wasting in adipose tissue [33,34]. An alternative explanation is that the production of hemocytes consumes significant metabolic resources, and when too high, restricts normal growth of the larva. To test this hypothesis, we measured triglyceride levels and visualized lipid droplets in the fat body. Interestingly, triglyceride contents decreased by 30%, 54% and 61% in fat bodies dissected from *eater^1^*, *pvf2* and *ras^V12^*-expressing larvae, respectively (Figures 1E, F). In contrast, *ras* depleted larvae with reduced hemocyte counts showed a 46% increase in triglyceride content compared to wild-type (Figure 1E). For all genotypes, triglyceride levels were inversely proportional to hemocyte counts. Fat body staining with Nile red and Bodipy confirmed triglyceride quantification (Figure 1G). Larvae with a higher number of hemocytes showed smaller lipid droplets of reduced intensity.

**Figure 1.**
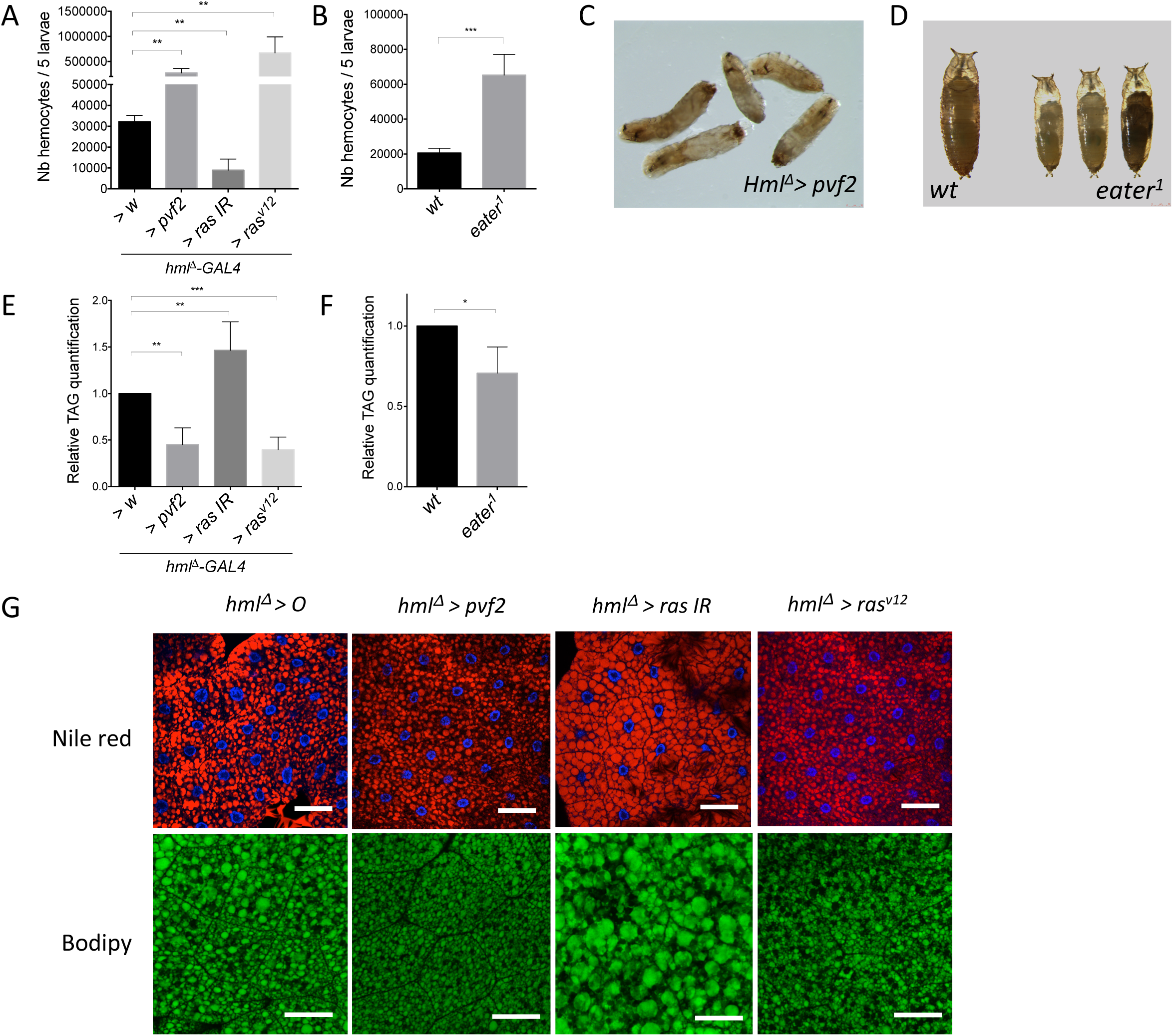
Fat body metabolism is linked to peripheral hemocyte proliferation. **(A, B)** Total counts of peripheral hemocytes from *hml^Δ^* > *w* (WT), *hml^Δ^*> *pvf2, hml^Δ^* > *ras* IR and *hml^Δ^* > *ras^v12^* **(A)** or WT (*hml^Δ^-GAL4,UAS-GFP*) and *eater^1^* mutant (*hml^Δ^-GAL4,UAS-GFP; eater^1^*) larvae **(B)**. Bars represent an average of 3-4 counts of 5 L3 wandering larvae with standard deviation (SD). **(C, D)** Representative images of dead *hml^Δ^*> *pvf2* **(C)** or WT and *eater^1^* mutant larvae **(D)** raised on poor diet. **(E, F)** Triacylglycerol (TAG) quantification of dissected fat bodies with the following genotypes: *hml^Δ^* > *w* (WT), *hml^Δ^*> *pvf2, hml^Δ^* > *ras* IR and *hml^Δ^* > *rasv12* **(E)** or wild-type (WT) (*hml^Δ^-GAL4,UAS-GFP*) and *eater^1^* (*hml^Δ^-GAL4,UAS-GFP;eater^1^*) **(F)**. TAG levels are normalized to corresponding protein levels. Graphs represent relative expression of an average of four measurements. **(G)** Representative confocal micrographs illustrating Nile red and Bodipy staining in fat bodies, corresponding to genotypes in (E). Scale bars correspond to 60 μm for Nile red and 33 μ

In contrast, *ras RNAi* larvae with a lower number of hemocytes exhibit larger lipid droplets and larger fat body cells. The fly CD36 homolog Croquemort (crq) has been shown to contribute to lipid-specific uptake in hemocytes [35]. If hemocytes and fat body compete for lipids, we expected that *crq^KO^* deficient larvae, that have wild-type number of hemocytes (Figure S2A), have higher fat store compared to wild-type. Indeed, Bodipy staining reveals larger lipid droplets of higher intensity in the fat body of *crq* deficient mutant compared to the wild-type (Figure S2B).

Collectively, these observations indicate that hemocyte production has a metabolic cost and that excessive hemocyte numbers affect larval viability upon nutrient deprivation, likely by depleting fat body energy storage.

### Nutrient availability influences peripheral hemocyte homeostasis

We next investigated whether the growth rate of the hemocyte compartment in larvae is fixed or whether it can be adjusted by nutritional cues. Considering the metabolic cost of hemocyte production and maintenance, it might be beneficial to reduce their proliferation in conditions of nutrient shortage to prioritize development of organs that are critical for survivals. In contrast, a rich diet might lead to an even higher number of hemocytes and ergo, a better immune defense. To test this hypothesis, we measured hemocyte numbers of third instar larvae raised on a poor diet consisting of only 20% nutrients compared to standard food or a high-fat diet obtained by supplementing regular food with lard (6%) as described in [35]. Strikingly, larvae raised on a poor diet had a 2.1-fold decrease in hemocyte count compared to larvae raised on a standard medium (Figure 2A). They also had fewer sessile hemocytes, no dorsal hemocyte patch and thinner lateral patches (Figure 2B). In contrast, larvae raised on a high-fat diet had a 1.5-fold increase in peripheral hemocyte numbers and showed more extended hemocyte patches on the dorsal side (Figure 2B, C). A higher number of hemocytes was also observed in larvae fed on a high glucose diet, indicating that this was not specific of the lipidic enriched food (Figure S2C). To confirm hemocytes variation according to food intake, we compared by RT-qPCR the size of the hemocyte compartment over the whole larva tissues by monitoring the expression of a hemocyte-specific gene, *hml*, and a ubiquitous gene (*rpL32)*. A decrease of the *hml/rpL32* ratio was observed when larvae were raised on a poor diet, consistent with a reduction of hemocyte number (Figure 2D). Of note, the *hml/rpL32* ratio was also lower under high-fat condition, suggesting that hemocyte compartment expands less compared to the whole larval body under rich food intake. These data point to the existence of a regulatory mechanism that couples blood cell proliferation to nutritive input.

**Figure 2.**
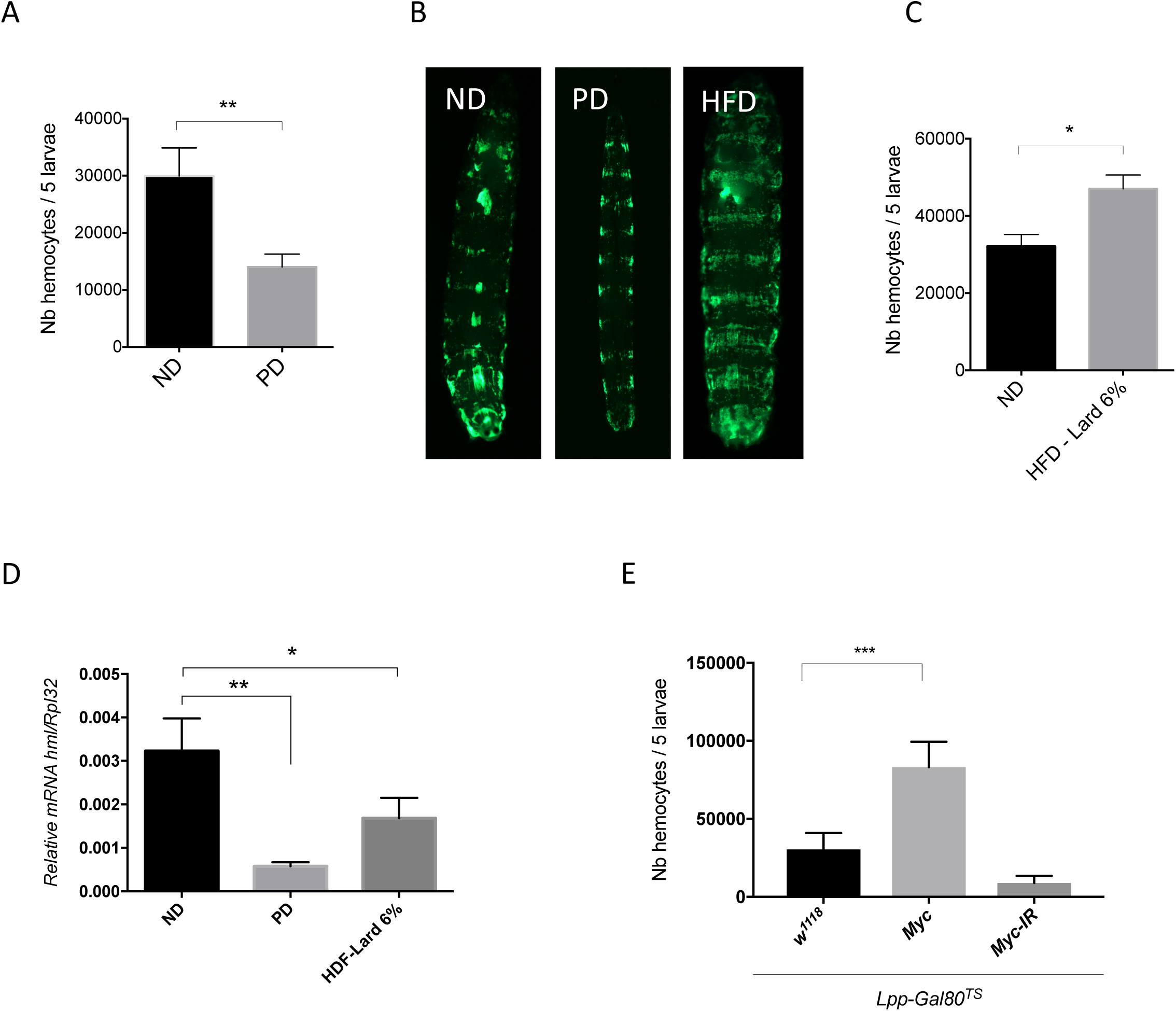
Nutrient availability influences peripheral hematopoiesis. **(A, C)** Total counts of peripheral hemocytes from *Hml^Δ^-GAL4,UAS-GFP* L3 wandering larvae fed on normal diet (ND), poor diet (PD) **(A)** or high fat diet with lard (HFD-lard6%) **(C)** from hatching. Bars represent an average of 3 counts of 5 L3 wandering larvae with SD. **(B)** Representative images illustrating *hml^Δ^-GAL4,UAS-GFP* L3 wandering larvae fed on normal diet (ND), poor diet (PD) or high fat diet (HFD). **(D)** RT-qPCR quantification of *hml* transcripts from fat bodies of mid-L3 larvae that were raised on ND, PD and HFD after hatching. **(E)** Total counts of peripheral hemocytes from *lpp^TS^ > w, lpp^TS^ > UAS-myc* and *lpp^TS^ > myc IR* at L3 wandering larval stage raised on ND. 10 ul hemolymph was loaded into a KOVA glasstic slide. Stars indicate P values of Student’s *t*-tests for **A** and **C** or Anova for **D** and **E**.

### Hemocyte number is influenced by the fat body metabolic state

The experiments described above indicate that hemocyte division and sessility are controlled by a factor that is sensitive to the metabolic state. In *Drosophila*, the fat body has been shown to sense and relay nutrient availability to coordinate the growth of the whole larva [36]. We hypothesized that hemocyte proliferation could be controlled by signals emanating from the larval fat body. To explore this hypothesis, we estimated the hemocyte number in larvae with reduced (*Lpp^TS^>Myc-IR*) or higher level of Myc (*Lpp^TS^>Myc*) in the fat body. As Myc is known to control organismal growth [37], we expected that a high level of Myc in the fat body could mimic the fed state, while lower Myc level would mimic the starved state. We observed that over-expression of *myc* in the fat body leads to a much higher hemocyte count while reducing Myc leads to a lower hemocyte number (Figure 2E). We conclude that the metabolic state of the fat body can influence the rate of peripheral hematopoiesis, suggesting the existence of signaling between these two tissues.

### *NimB5* is expressed in the fat body upon nutrient scarcity

We then searched for genes expressed in the fat body that encode secreted factors susceptible to regulate hemocyte number according to the metabolic state. Our attention focused on *NimBs*, which encode secreted proteins of the Nimrod family; some of them being expressed in the fat body at the larval stage. The Nimrod family comprises six transmembrane proteins (NimA; NimC1, C2, and C4 (SIMU); Draper and Eater) and five secreted proteins (Nimrod B (NimB) family: B1, B2, B3, B4, and B5) with characteristic EGF-like repeats, also called “NIM repeats”[38]. Studies have revealed the implication of the transmembrane Nimrod proteins in bacterial phagocytosis (NimC1, Eater) and apoptotic corpses (SIMU, Draper), and in hemocyte sessility and adhesion (Eater) [30,39–43]. So far, none of the secreted NimBs have been functionally characterized. Interestingly, one *NimB*, *NimB5*, has been shown to be induced in a mitochondrial mutant, that mimics a starvation state [44]. To better characterize *NimB* expression profiles, we monitored their transcripts in larvae raised 4h, 8h, and 16h on a poor diet from early L3 stage. *NimB5* was the only gene significantly upregulated with a three-fold induction level compared to wild-type at 8h (Figure 3A). Another experiment confirmed that the level of *NimB5* expression was increased when larvae feed on agar medium (containing only agar and water) (Figure S2D).

**Figure 3.**
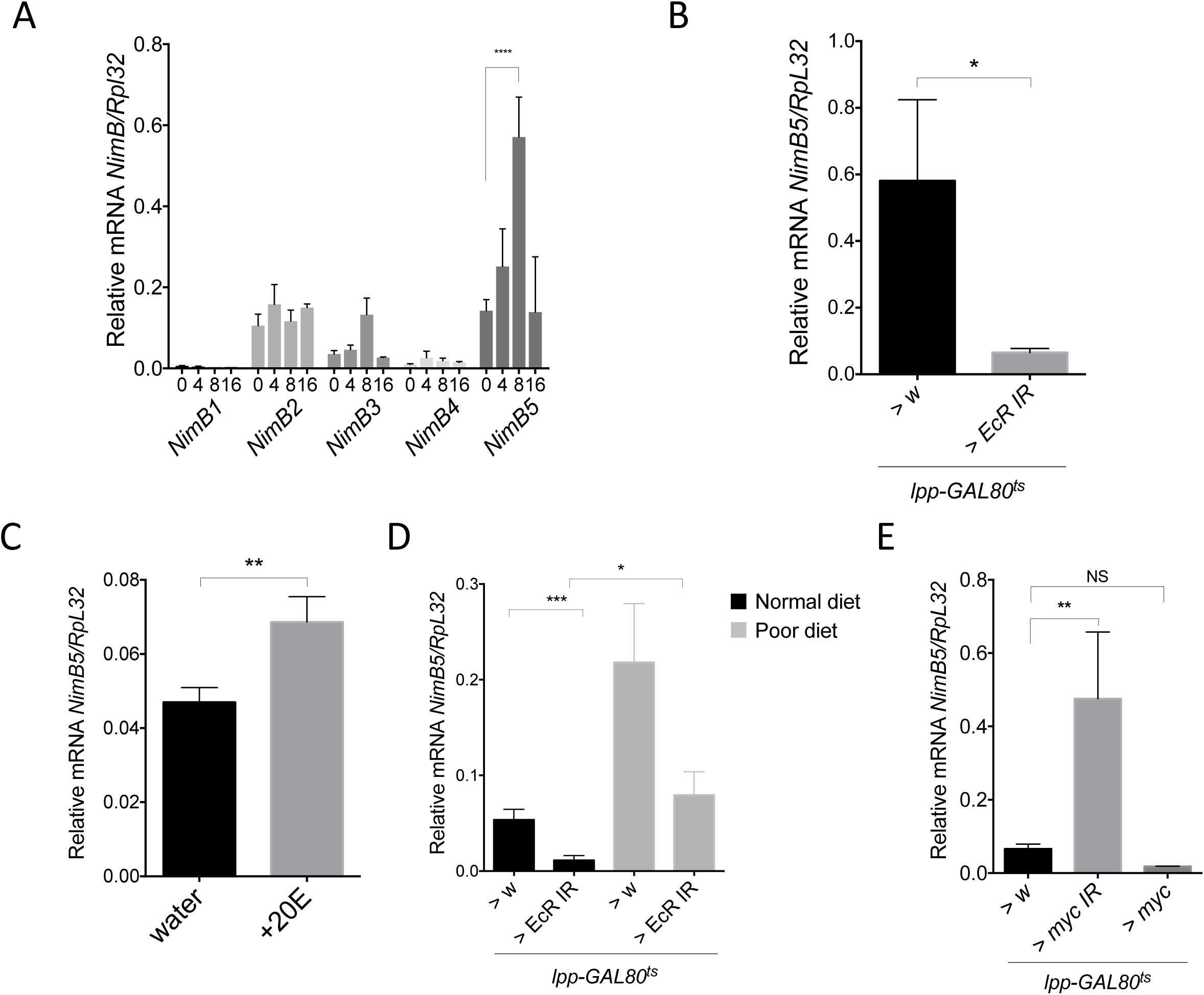
NimB5 expression is activated upon starvation and controlled by ecdysone. **(A)** RT-qPCR quantification of *NimB1, NimB2, NimB3, NimB4* and *NimB5* transcripts from fat bodies of mid-L3 larvae raised on poor diet dissected at indicated time points. Bars represent an average of three measurements with 15 animals for each experiment. **(B)** *NimB5* transcripts from fat bodies of *lpp^TS^ > w* and *lpp^TS^ > Ecd IR* L3 wandering larvae. **(C)** *NimB5* transcripts from fat bodies of mid-L3 larvae raised for 4h on crushed banana supplemented with water or 20-hydroxyecdysone (final concentration 0.5 mM). **(D)** *NimB5* transcripts from fat bodies of animals raised on normal or poor diet, after 8h of *EcR* silencing in fat bodies **(E)** Relative expression of *NimB5* to *RpL32* mRNA levels in fat bodies of *lpp^TS^ > w, lpp^TS^ > myc IR* and *lpp^TS^ > UAS-myc* from L3 wandering larvae. Bar graph data are presented relative to *RpL32* as mean ±SEM. Graphs represent an average of 3 measurements with 15 animals for each experiment. Stars indicate P values of Anova for **A, D** and **E**, or Student’s *t-*test for **B** and **C.**

Interestingly, published ModENCODE developmental and tissue array data showed that *NimB5* is almost exclusively expressed in the fat body of larvae [45], with the expression peaking at the wandering stage when larvae migrate away from food and prepare for pupariation. This developmentally induced starvation precedes the extended pupal feeding arrest. The RNA profiles matched proteomic profiles performed on whole animals that confirmed the increased *NimB5* expression at the onset of pupariation and higher levels as animals reach the end of the pupal stage and emerge [46].

### *NimB5* is under the control of the growth metabolic regulators EcR and Myc

The rapid and high increase of *NimB5* expression during the late larval stage, as well as its sustained expression during the pupal stage, coincides with the peak of ecdysone titers at the larval-pupal transition and the high titers in pupae, respectively. Therefore, we checked whether the developmental expression of *NimB5* could be controlled by ecdysone. For this, we silenced the gene encoding the ecdysone receptor (EcR) specifically in the fat body (*lpp^TS^>EcR-RNAi*) at the mid-L3 larval stage to limit the impact on animal growth [47]. We observed lower *NimB5* transcript levels in *lpp^TS^>EcR-RNAi* wandering L3 larvae compared to wild-type (Figure 3B). Consistent with the role of ecdysone in the regulation of *NimB5*, feeding mid-L3 larvae for five hours with the steroid hormone 20-hydroxyecdysone significantly increased *NimB5* expression in the fat body (Figure 3C). However, the *NimB5* gene was still inducible to some extent by starvation in larvae with reduced fat body EcR signaling (Figure 3D). This result indicates the existence of other signaling inputs to regulate *NimB5* gene expression upon nutrient scarcity.

The transcription factor Myc is a well-established downstream target of EcR signaling in the fat body [47]. Myc is repressed upon EcR activation and is known to control organism growth. Thus, we hypothesized that *NimB5* could be negatively regulated by Myc, as *NimB5* is expressed in conditions of reduced growth. Accordingly, silencing *Myc* specifically in the fat body significantly increased *NimB5* expression, whereas overexpressing *Myc* in the fat body led to a slight downregulation of *NimB5* without reaching significance (Figure 3E). We conclude that *NimB5* regulation receives input from both Myc and EcR, two key metabolic integrators of the fat body.

### NimB5 binds on hemocytes

NimB5 belongs to the Nimrod family that has been shown to regulate many essential biological functions of hemocytes [38]. This function raised the hypothesis that NimB5 produced by the fat body could bind to hemocytes. To test this notion, we generated transgenic fly lines carrying a *UAS*-*NimB5-RFP* or an endogenously GFP-tagged *NimB5* locus fusion (derived from the Dresden pFlyfos collection) [48]. Overexpression of the *NimB5-RFP* fusion protein in the fat body with *lpp^TS^-Gal4* in L3 larvae confirmed that NimB5 is indeed secreted, as RFP fluorescent signals were observed in nephrocytes, which recycle hemolymph proteins (Figures S3A, B). Strikingly, hemocytes collected from these larvae exhibited small RFP dots on their surface, showing that NimB5-RFP does indeed bind to circulating hemocytes (Figure 4A). Immunostaining using flies carrying the endogenously *GFP*-tagged *NimB5* locus confirm that NimB5 localizes to the surface of hemocytes (Figure 4B, C). Histological sections revealed the presence of NimB5-RFP signals on the membrane of sessile hemocytes (Figure 4D, white arrows).

**Figure 4.**
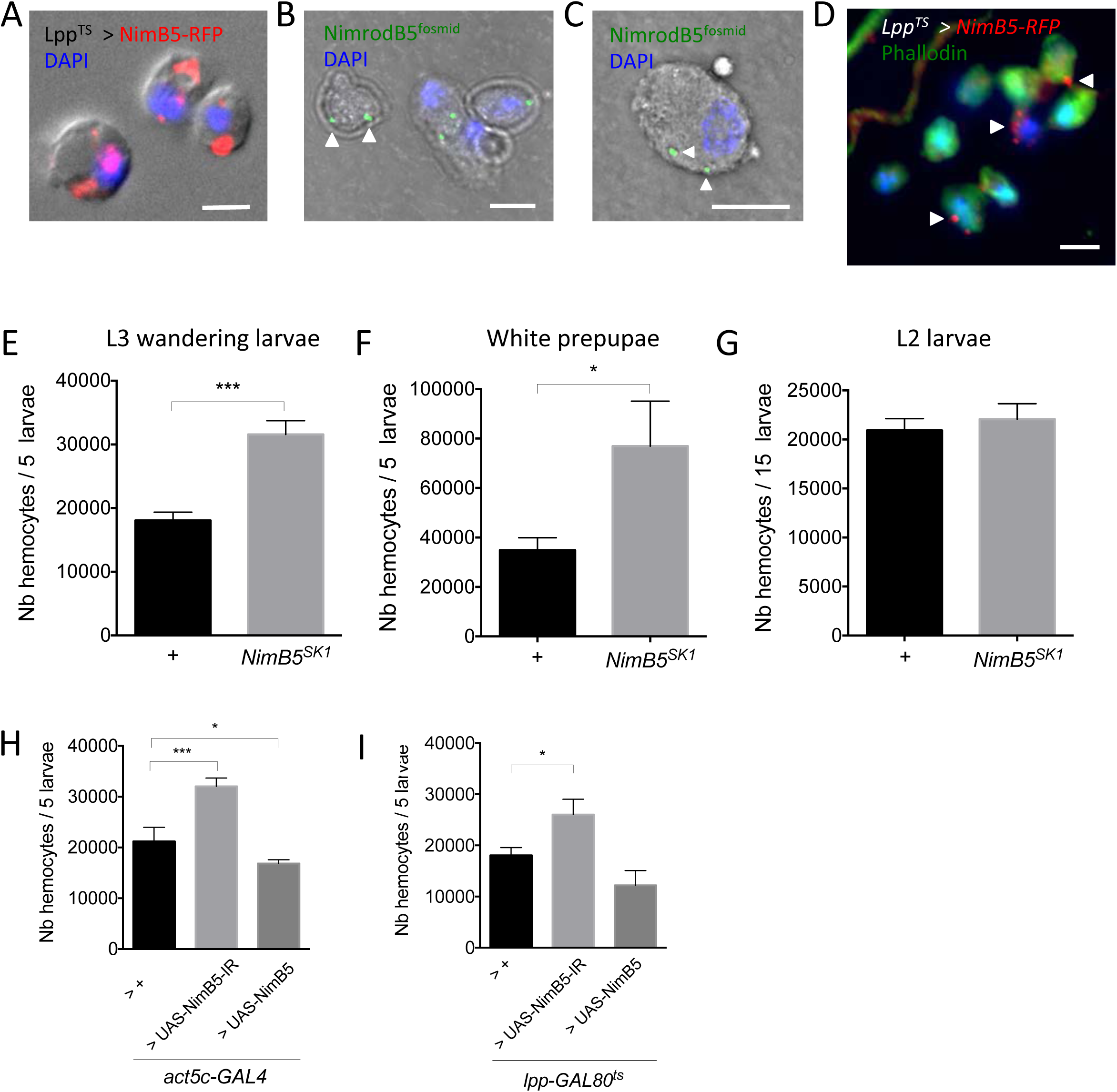
NimB5 binds to blood cells and controls hemocyte number. **(A)** Representative image of hemocytes stained with DAPI (blue), illustrating the localization of NimB5-RFP (red) secreted from fat body (*lpp^TS^* > *UAS-NimB5-RFP*). Overlay of fluorescence and DIC. Scale bar = 10 μm. **(B, C)** Confocal microscopy images of hemocytes stained with DAPI, illustrating the localization of endogenous NimrodB5. Hemocytes are extracted from larvae carrying Flyfos-*NimB5* transgene and stained against V5 antigen. Overlay of fluorescence and phase contrast images. Scale bar = 10 μm. **(D)** Cross sections of L3 wandering larvae, illustrating hemocyte patches with NimB5-RFP secreted from fat body (*lpp^TS^* > *UAS-NimB5-RFP*). Scale bar = 10 m. **(E, F, G)** Peripheral (sessile and circulating) hemocyte counts of *w^1118^* or *NimB5^SK1^* animals combined with *hml^Δ^*.*dsred.nls* from L3 wandering larvae (**E**), white prepupae (**F**) and L2 larvae (**G**). **(H)** Peripheral hemocyte counts of *act5c > w, act5c > nimB5-IR* and *act5c > UAS-NimB5* larvae. **(I)** Peripheral hemocyte counts of *lpp^TS^ > w*, *lpp^TS^ > NimB5-IR*; *lpp^TS^ > UAS*-*NimB5*. Results represent a total of five animals from three individual experiments. We applied Student’s *t*-tests for **E, F, G** or Anova test for **H** and **I**. LG = lymph gland.

### NimB5 negatively regulates hemocyte numbers

The inducibility of *NimB5* upon nutrient scarcity and its ability to bind to hemocytes makes it an excellent candidate to contribute to a fat-body hemocyte signaling. To further investigate the function of NimB5 in peripheral hematopoiesis, we generated a null mutation in the *NimB5* gene by Crispr-Cas9, referred to as *NimB5^SK1^*. *NimB5^SK1^* has a 13bp deletion in the first exon, inducing a premature stop codon and leading to a truncated protein of 54 amino acids instead of 351 (Figures S4A, B). *NimB5* mutants were viable and did not show any morphological defect. We then investigated whether the *NimB5* mutation affects hemocyte numbers by counting both sessile and circulating hemocytes using a method developed by Petraki et al., [49] coupled with flow cytometry. *NimB5^SK1^* third instar larvae and pupae had, respectively, 1.7 and 2.2 times more hemocytes compared to wild-type when raised on a standard diet (Figures 3E, F). These higher hemocyte numbers in *NimB5^SK1^* larvae were not due to higher numbers of embryonic hemocytes since *NimB5^SK1^* and wild type larvae contained comparable numbers at the L2 stage (Figure 4G). Since hemocyte numbers markedly vary from one genetic background to another, we performed additional experiments to confirm that the *NimB5* mutation is responsible for a higher number of hemocytes.

First, we introgressed the *NimB5^SK1^* mutation by successive backcrosses into the *w, Drosdel* background [50]. Figure S5A shows that isogenized *NimB5^SK1^* larvae also have higher hemocyte numbers compared to their *w, Drosdel* counterparts. Second, higher hemocyte numbers were also observed in trans-heterozygous *NimB5^sk1^*/*Df(2L)BSC252* larvae carrying the mutation over a deficiency that removes the *Nimrod* locus (Figure S5B). We noticed that *NimB5^sk1^*/*Df(2L)BSC252* larvae, however, had fewer hemocytes compared to homozygous *NimB5^SK1^* larvae. This result is likely because the deficiency removes several other *Nimrod* genes, which may also influence peripheral hematopoiesis.

Furthermore, the ubiquitous *RNAi*-mediated knockdown of *NimB5* using the *actin-GAL4* driver led to a significant increase in the hemocyte population (Figure 4H). To dissect in which organ/tissue NimB5 was required, we used fat body and hemocyte-specific drivers. Silencing *NimB5* in the larval fat body (using *lpp^TS^-Gal4* or *C564-Gal4*) but not in hemocytes (*hmlΔ-Gal4*) also resulted in higher hemocyte numbers compared to sibling larvae expressing only the Gal4 construct (Figures 4I; S5C, D). Consistent with its role as a negative regulator of hemocyte proliferation, overexpression of *NimB5* using *actin-Gal4* led to significantly lower hemocyte numbers (Figure 4H). Overexpressing *NimB5* in the fat body with either *lpp^TS^-Gal4* or *C564-Gal4* also led to reduced hemocyte numbers, although the difference did not reach statistical significance (Figures 4I and S5C). These discrepancies could be explained by the stronger expression of the ubiquitous *actin-Gal4* driver compared to fat body-specific Gal4 constructs, which would result in higher levels of NimB5 in the hemolymph. Overexpression of *NimB5* using the *hmlΔ-Gal4* driver in an otherwise wild-type background did not affect hemocyte counts (Figure S5D). Nevertheless, overexpressing *NimB5* in the fat body or hemocytes of *NimB5^SK1^* mutant larvae rescued the over-proliferation phenotype by reducing hemocyte numbers to normal levels (Figure S5E, F).

Consistent with the increased hemocyte counts, *NimB5^SK1^* mutants showed larger sessile hemocyte patches beneath the cuticle, but also more circulating hemocytes (Figure 5A). Phosphohistone 3 (PH3) staining and EdU incorporation assays [19] showed that *NimB5^SK1^* larvae exhibited a higher frequency of mitotic hemocytes compared to wild-type (Figures 5B, C). Histological analysis and EdU incorporation experiments showed that the *NimB5^SK1^* mutation did not affect lymph gland hematopoiesis (Figures 5D, E). Importantly, hemocytes in the *NimB5^SK1^* mutants appeared fully functional since they were perfectly able to phagocytose bacteria (Figure S6A). The mutant larvae did not show any significant defect in melanization or encapsulation, indicating that *NimB5^SK1^* larvae retain the ability to differentiate crystal cells and lamellocytes (Figures S6B-E). The absolute number of crystal cells per larva was higher in *NimB5^SK1^* larvae compared to the wild-type as expected for a mutant with higher hemocyte counts (data not shown), however, the ratio of crystal cells over plasmatocytes was lower (Figure S6F). Collectively, our data reveal that a secreted factor emanating from the fat body upon nutrient scarcity, NimB5, negatively regulates the rate of peripheral hemocyte proliferation without impeding hemocyte function.

**Figure 5.**
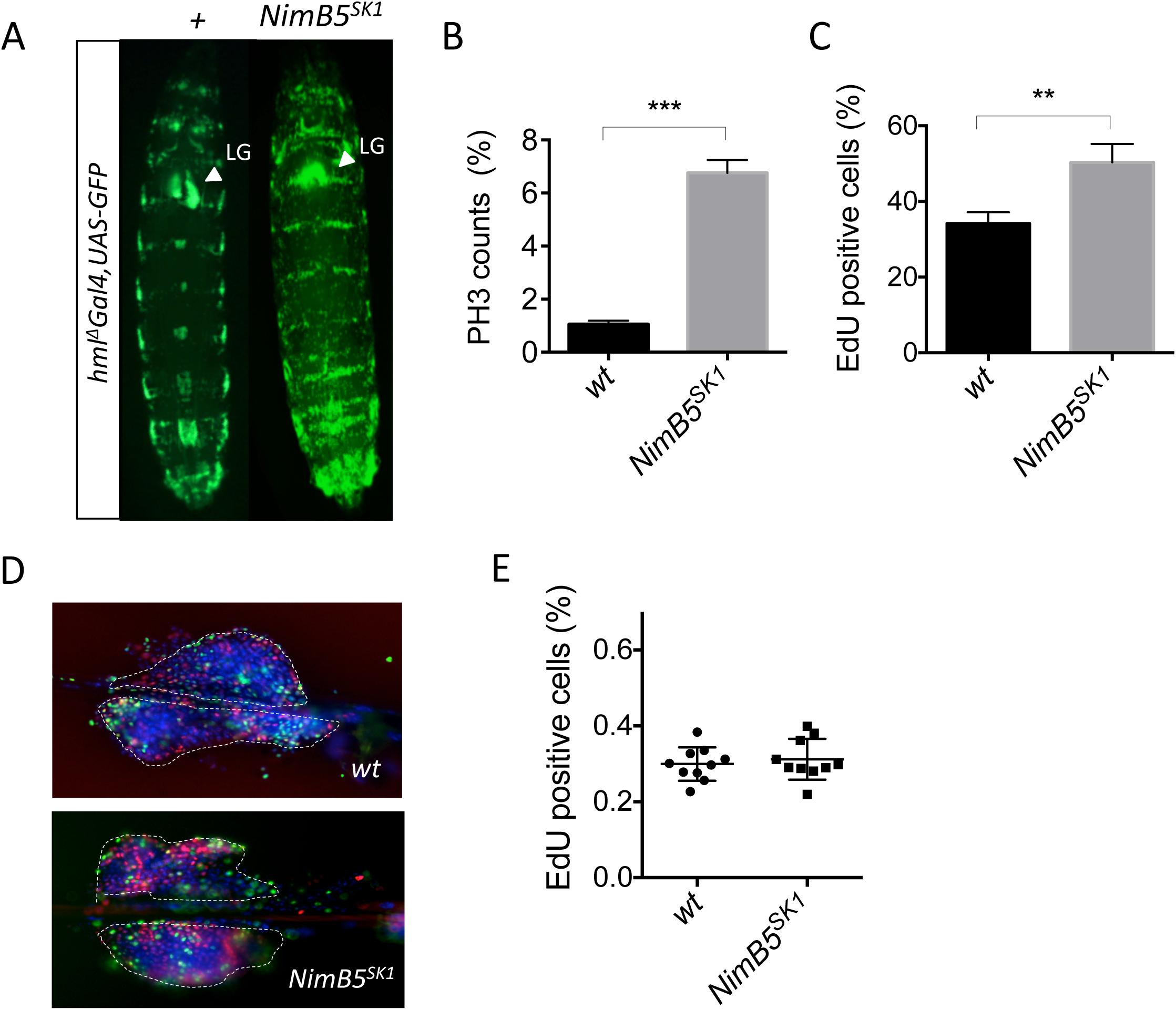
NimB5 blocks hemocyte proliferation. (A) Images of live wt *hml^Δ^-Gal4,UAS-GFP* and *NimB5^SK1^,hml^Δ^-Gal4,UAS-GFP* L3 wandering larvae. Percentages of PH3 **(B)** or EdU **(C)** positive cells in peripheral cells positive for *hml^Δ^.dsred.nls*, from *w^1118^* and *NimB5^SK1^* at L3 wandering stage. **(D)** Images of lymph glands from *w^1118^* and *NimB5^SK1^* genotypes combined to *hml^Δ^.dsred.nls* marker (red) from L3 wandering larvae, stained with EdU proliferation marker (green). **(E)** Percentages of EdU positive cells from 10 lymph glands of *w^1118^* or *NimB5^SK1^* animals combined with *hml^Δ^.dsred.nls* marker at L3 wandering larval stage. For **B, C** *P* values from Student’s *t-*test.

### NimB5 affects hemocyte sessility and adhesion

Adherent cells, notably when establishing contacts with other cells, are less proliferative, a process called “contact inhibition of proliferation” [51]. This raised the question of whether hemocyte proliferation could be associated with decreased adhesion, or hemocytes establishing fewer contacts with neighboring cells. We thus explored whether NimB5 affects hemocyte adhesion and sessility. First, we seeded wild-type or *NimB5^SK1^* hemocytes on glass, allowed them to adhere for 30 min, then performed phalloidin staining. Image analysis revealed that hemocytes of *NimB5^SK1^* mutants spread much less compared to wild-type (Figures 6A, B). This spreading defect was not due to smaller hemocyte size, as revealed by a Tali*™* Image-Based Cytometer (Figure S7).

**Figure 6.**
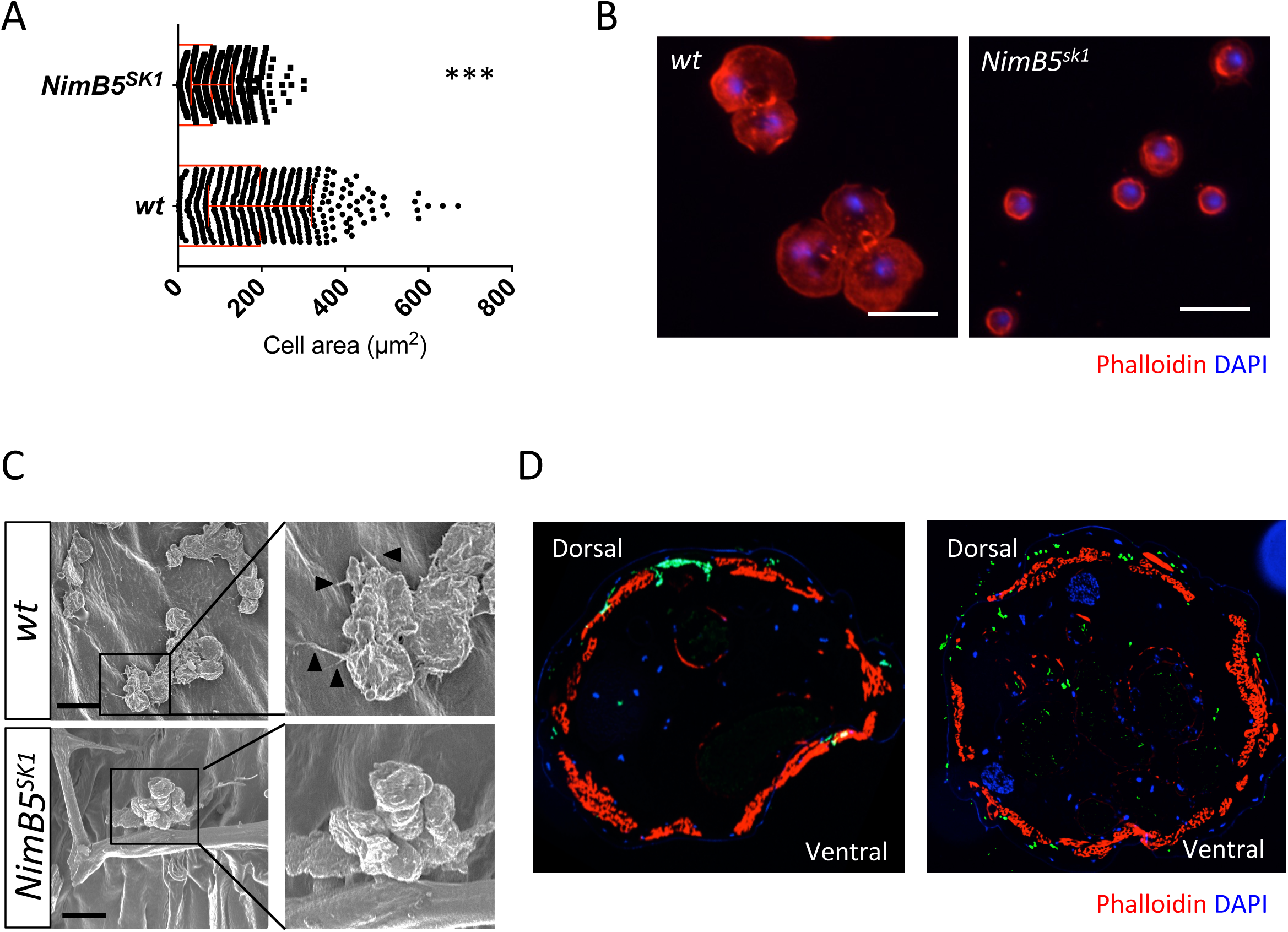
NimB5 promotes hemocyte adhesion *in vitro* and *in vivo*. **(A)** Bar diagram of hemocyte areas from *w^1118^* and *NimB5^SK1^* larvae at L3 wandering stage. Hemocytes were allowed to spread on slides for 30 min and were stained with AF488 phalloidin. **(B)** Representative images of hemocytes collected from *w^1118^* and *NimB5^SK1^* larvae at L3 wandering stage, stained with rhodamine phalloidin. Scale bar = 10 m. **(C)** Scanning electron microscopy of internal cuticle side of L3 wandering larvae. *w^1118^* and *NimB5^SK1^* animals were opened ventrally and all organs were removed to allow visualization. Scale bar = μm. **(D)** Cross-sections from wild-type and *NimB5^SK1^* larvae with *hml*Δ-*Gal4,UAS-GFP* at L3 wandering stage. Phalloidin staining in red, DAPI in blue. For **A** *P* values from Mann-Whitney test.

To assess how NimB5 affected hemocyte sessility *in vivo*, we subsequently performed scanning electron microscopy analysis of sessile hemocytes, focusing on the inner side of the cuticle of L3 wandering larvae. We observed that hemocytes in the *NimB5^SK1^* mutant spread less and lacked filopodia as compared to wild-type (Figure 6C, black arrows). Finally, we recombined the *hml^Δ^-Gal4,UAS-GFP* hemocyte marker with the *NimB5^SK1^* mutation to analyze the peripheral hemocyte localization. Although *NimB5^SK1^* mutant larvae still contained sessile hemocyte patches, the cells were only loosely attached to the inner wall of the larvae (Figure 6D). We next asked whether *NimB5^SK1^* larvae also have defects in the other sessile hemocyte type, the crystal cell [17]. Sessile crystal cells derived plasmatocyte by a process of transdifferentiation from sessile plasmatocytes [30,52]. Thus, we expected that a defect in plasmatocyte sessility in *NimB5^SK1^*mutant larvae should affect the number of crystal cells. Heating larvae in water for 30 min at 67°C causes spontaneous activation of the prophenoloxidase zymogen within crystal cells and their subsequent blackening, making them sessile crystal cells visible through the cuticle as black puncta [53]. Consistent with a role of NimB5 in sessility, *NimB5^SK1^* mutant showed reduced black puncta, which is likely a secondary consequence of a defect in plasmatocyte sessility (Figures S6G, H, see below). We conclude that the fat body factor NimB5 regulates not only proliferation but also hemocyte adhesion properties and sessility.

### NimB5 contributes to the survival of larvae raised on a poor diet

Our data show that NimB5 is a fat body derived factor, upregulated in response to nutrient scarcity, and required to maintain physiological hemocyte numbers. We hypothesized that if NimB5 regulates hemocyte numbers in response to nutrient availability, *NimB5* mutants should exhibit fitness defects when raised on a poor diet. Thus, we shifted *NimB5* and wild-type mid-instar larvae from a rich to a poor diet and analyzed adult eclosion rate, and indeed, most *NimB5* but not wild-type animals showed significant pupal lethality at the pharate stage (Figures 7A, B). We speculated that this lethality was due to high hemocyte numbers since *NimB5* mutants were unable to fully down-regulate hemocyte proliferation under poor diet conditions as the wild-type did (Figure 7C). To demonstrate that the lethality was indeed linked to an excess of hemocytes, we eliminated them in *NimB5^SK1^* mutants by overexpressing the pro-apoptotic *bax* gene using the *hml^Δ^-GAL4* driver. Previous reports have demonstrated that hemocyte-depleted (hemoless) *Drosophila* have an almost normal development [54,55]. Expression of *bax* in hemocytes of *NimB5* deficient larvae suppressed the pupal lethality observed in poor diet conditions (Figure 7D). We then explored the amount of lipid in the fat body of *NimB5* deficient larvae with or without hemocytes. As expected for a higher hemocyte count mutant, Bodipy stainings show that *NimB5* larvae have smaller lipid droplets compared to the wild-type (Figure 7E). In contrast, *NimB5* hemoless larvae expressing the pro-apoptotic gene in hemocytes have bigger lipid droplet confirming that hemocytes affect lipid store. Taken together, these results demonstrate that fat body-hemocyte regulatory loop which involves NimB5 is critical for survival on a poor medium.

**Figure 7.**
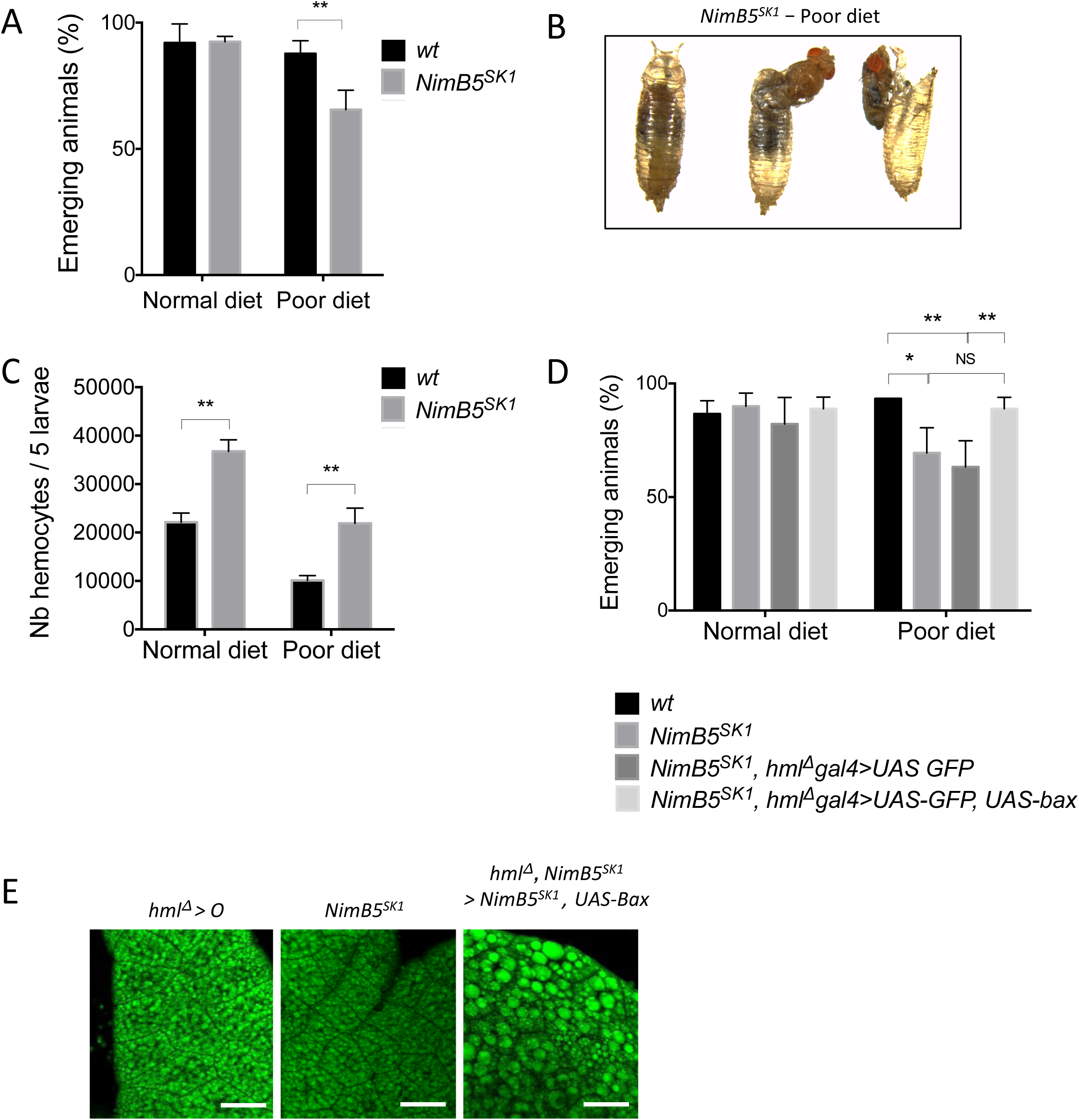
NimB5 promotes adaptation to poor diet. **(A)** Percentages of emerging *WT* and *NimB5^SK1^* animals raised on normal diet until mid-L3 stage and then transferred to fresh normal diet or poor diet. **(B)** Representative image of non-emerging *NimB5^SK1^* animals. Development stops at pharate stage. **(C)** Peripheral hemocyte counts from *w^1118^* or *NimB5^SK1^* larvae combined to *hml^Δ^.dsred.nls* marker in L3 wandering stage. Animals were raised on normal diet until mid-L3 stage and transferred to new vials with normal diet or poor diet until L3 wandering larval stage. **(D**) Percentages of emerging *WT* and *NimB5^SK1^* animals expressing the pro-apoptotic gene *bax* specifically in hemocytes (*hmlΔ-Gal4,UAS-GFP* > *UAS-bax).* Animals were raised on normal diet until mid-L3 stage and transferred to new vials with normal diet or poor diet. Experiments were done with 25 animals for each genotype, at 25°C for A and D. Results represent the average of three experiments. (**E**) Representative confocal micrographs illustrating Bodipy staining in fat bodies from *hml^Δ^* > *w*, *nimB5^SK1^* and *hmlΔ-Gal4,UAS-GFP* > *UAS-bax* combined with *nimB5* mutation. Scale bars correspond to 33 μm for Bodipy. For **A, C** and **D**, *P* values from Anova test.

## Discussion

Nutrition is an essential environmental parameter in the determination of body size. In Metazoans, the coordination of growth among different body parts according to environmental conditions relies on cross-talk between different organs via the production of long-range signaling molecules [56]. In this study, we uncover an adipokine, NimB5, which is produced by the fat body upon nutrient scarcity to reduce hemocyte adhesion and proliferation (See model on Figure S10). The disruption of this signaling loop results in conditional lethality when larvae are raised on a poor diet. Our study shows that larvae die from their inability to reduce hemocyte numbers, which is likely detrimental to energy storage and not from an over-activation of the immune response. In this article, we have privileged the notion that the depletion of the metabolic store due to the high energy demand of numerous hemocytes causes larval lethality on a poor diet. This is supported by the observation that lethality on poor diet correlates with hemocyte number. At first sight, it is surprising that a two-threefold increase in hemocyte number is so detrimental to animal fitness, taking into consideration the small size of the hemocyte compartment compared to other tissues. It is likely that *Drosophila* plasmatocytes are strong energy users, as mammalian macrophages, and energetically costly. Nevertheless, we cannot definitely exclude that the lethality we observed has another cause. For instance, this lethality could be due to a hemocyte factor that directly regulates fat body metabolism rather than metabolic competition between tissues. Further studies are required to better characterize the interactions between the fat body and hemocytes.

### Adjustment of hemocyte numbers to the nutritional status

Our study demonstrates that the growth of the larval hemocyte compartment is not stereotypical but influenced by nutritional cues. Larvae fed on a rich diet have higher hemocyte numbers, while larvae experiencing nutrient deprivation have reduced hemocyte counts. This reveals flexibility in the way an organism allocates energy to build the immune system. We further show that this adjustment is critical since hemocyte production has a significant metabolic cost and can compete with the fat body building up metabolic stores. Consequently, larvae with higher hemocyte numbers fail to develop correctly when raised on a poor diet. Even when grown on a rich diet, these larvae store lower lipid amounts in their fat body. This indicates that a high hemocyte population has a critical metabolic cost, which cannot be fully compensated through increased nutrient uptake. The adjustment of hemocyte numbers to nutritional cues might explain the high variability in hemocyte numbers observed both within and between fly stocks among laboratories using different fly mediums and distinct crowding conditions. While the coordination of organ growth is the focus of intense studies in *Drosophila* research, the variation in the size of blood cell compartments is rarely taken into consideration. As hemocytes are mainly dispensable in normal conditions [54,55], the reduction of their numbers offers a mechanism to survive adverse dietary conditions. Nevertheless, this adaptation comes at the expense of a robust immune system. The existence of regulatory mechanisms tailoring investment in the immune system to nutrient availability is likely to be a general principle of animal host defense, to reduce potential trade-offs between immunity and other physiological functions.

In 1997, a landmark paper demonstrated that *Drosophila* selected for improved resistance against a parasitoid wasp had reduced larval competitive ability, notably when raised on poor diet, pointing to the existence of a trade-off between a robust immune system and growth [7]. Later on, effective resistance to parasitoid wasps in selected lines was associated with a doubling of hemocyte numbers [57,58]. These studies are entirely consistent with our observation that increased size of the hemocyte compartment affects the ability of larvae to survive on a poor diet. Our present study sheds some light on the molecular mechanisms underlying this trade-off, showing that excessive hemocyte numbers hamper fat body energy storage. The existence of phenotypic plasticity in the growth of peripheral hematopoiesis is likely adaptive for fruit flies that live on rotting fruit, an ephemeral ecosystem, and frequently experience nutrient stress. As the maintenance of hemocytes tends to affect metabolic storage, we cannot exclude that the increased resistance to parasitoids of larvae with higher hemocyte counts does not result from more potent cellular immunity, but is instead due to the depletion of lipid storage which is a critical resource for parasitoids [58,59]. In this line, we performed survival analyses, which revealed that endoparasitoids have a lower success when they infest *NimB5* mutants, compared to wild-type, due to increased pupal lethality in infested mutants (Figure S8). This observation could likely be explained by the higher hemocyte number leading to a stronger encapsulation reaction or the depletion of lipids that prevents the development of both the *Drosophila* larvae and the wasps.

### Regulation of NimB5 by central metabolic cues

We demonstrate that NimB5 functions as an adipokine to limit hemocyte proliferation. Accordingly, the expression of the gene is upregulated at the third instar wandering stage, when larvae stop feeding before entering pupariation. *NimB5* is also induced by nutrient shortage mainly at early time points, which suggests that it is involved in the early adaptation to poor food. During development, the expression profile of *NimB5* is very similar to that of *Dilp6*, which encodes an insulin-like peptide produced by the fat body to regulate growth at non-feeding stages [60]. *Dilp6* is also induced by starvation. Much like *Dilp6*, the Ecdysone Receptor positively regulates *NimB5* expression. In *Drosophila*, ecdysone is not only involved in the activation of molts and metamorphosis but also plays a growth-inhibitory function during larval development, notably by inhibiting Myc in the fat body [47,61]. Thus, *NimB5* is regulated in the fat body by key metabolic pathway integrators which signal ‘growth arrest’, as expected for a gene encoding an adipokine that informs hemocytes to stop proliferating. Since *NimB5* is still induced in *EcR* deficient larvae upon starvation, this gene likely receives additional regulatory inputs.

Previous studies have shown that nutritional deprivation also impinges on the maintenance of blood progenitors in the lymph gland causing the expansion of mature blood cells. These studies reveal that both systemic and local signals (e.g. TOR and insulin pathways) regulate blood progenitor maintenance in the lymph gland [62–66]. The observation that NimB5 does not regulate hematopoiesis in the lymph gland reveals that distinct mechanisms upon nutrient deprivation regulate central (lymph gland) and peripheral hematopoiesis. The existence of NimB5 likely provides more versatility in the control of animal growth by uncoupling hemocyte development from that of other organs, which is regulated by more generic growth signals (e.g. insulin-like peptides).

### NimB5 promotes adhesion and inhibits proliferation

Our study reveals that NimB5 binds to peripheral hemocytes to regulate adhesion and proliferation. In this article, we have favored the notion that increased hemocyte proliferation in *NimB5* larvae is a secondary consequence of an adhesion defect, which prevents cells from interacting with each other. Future studies should address whether this proliferation defect is a consequence of impaired adhesion, or whether adhesion and proliferation are uncoupled. An important point is to clarify whether NimB5 function is a growth factor that activates a downstream signaling pathway in hemocytes and/or directly favors hemocyte-hemocyte contacts. Of note, NimB5 could inhibit hemocyte adhesion, preventing sessile hemocytes from being under the influence of the neuronal system, as shown by Makhijani et al. [19,67]. The phenotype of *NimB5* animals shared many similarities with *eater*, suggesting that the secreted NimB protein and the transmembrane receptor Eater, could function in the same process. The observation that *eater^1^, NimB5* double mutants exhibit even higher hemocyte counts (Figure S9) than single mutants suggests that NimB5 could also interact with another receptor. *eater* null mutants show a stronger hemocyte phenotype than *NimB5* mutants both in term of hemocyte increase and defect in sessility. This could also be explained by the implication of other NimB members. The involvement of other NimB is also supported by the observation that *NimB5* deficient larvae have the ability, albeit reduced compared to the wild-type, to limit hemocyte population expansion when raised on a poor diet (Figure 7C). The Nimrod gene family has experienced diversification in *Drosophila* species. We hypothesize that the expansion of the Nimrod family, with five secreted forms and six proteins bearing a transmembrane domain, could underlie the constitution of a complex platform to regulate peripheral hematopoiesis, notably hemocyte proliferation and sessility. Future studies should characterize the role of the other secreted Nimrod members, notably those expressed in the fat body.

## Materials and methods

### Drosophila stocks and rearing conditions

All fly stocks used in this work are defined in the supplementary materials table and references therein. All stocks were reared on standard fly medium comprising 6% cornmeal, 6% yeast, 0.62% agar, 0.1% fruit juice, supplemented with 10.6g/L moldex and 4.9ml/L propionic acid. Fly stocks and crosses were maintained at 25°C on a 12 h light/ 12 h dark cycle. For poor diet experiments, mid-L3 larvae were raised on medium that corresponds to 20% nutrients of regular medium, with 1.2% cornmeal, 1.2% yeast, 0.62% agar supplemented with 0.98ml/L propionic acid. L2 larvae were selected 48-52 hours after egg laying (AEL), early L3 larvae were selected 72-90 hours AEL (3 days), mid-L3 larvae were selected 96-110 hours AEL (4 days) and L3 wandering larvae were selected 110-120 hours AEL (5 days).

### Plasmids and transgenic lines

For overexpression and complementation studies, the genomic region from the 5’UTR to the stop codon of the *NimB5* gene was amplified from BACR09N24 and cloned into the pDONR207 Gateway vector (Invitrogen) and subsequently sub-cloned in the pTW (Drosophila Genomics Resource Center plasmid) transgenesis vector and used to generate transgenic *UAS-NimB5* flies. For the *UAS-NimB5-RFP*, the *NimB5* cDNA sequence without STOP codon was cloned into the entry vector pENTR/D-Topo (Invitrogen) and subsequently shuttled into the RFP expression vectors pTWR (C-terminal RFP tag), obtained from the DGRC *Drosophila* Gateway vector collection. Plasmids were injected either at the Fly facility platform of Clermont-Ferrand (France) or by BestGene Inc. (Chino Hills, CA, USA).

### Hemocyte counts by flow cytometry

Hemocytes were bled into 120 µL of PBS containing 1 nM phenylthiourea (PTU, Sigma) to prevent melanization. Hemocytes were counted based on their morphometric properties with a BD Accuri C6 flow cytometer, USA. Cells were selected based on their size with a FSC-A from 2.5.10^6^ to 3.5.10^6^ and based on their granularity with a SSC-A from 1.8.10^5^ to 2.5.10^5^. In a second part, cells were gated for singlets (FSC-H versus FSC-A). For fluorescent hemocytes, we used FL1 (GFP) or FL2 (dsred) detectors.

### Triglyceride measurements

We quantified free glycerol and all CCA substrates with the free glycerol reagent (F6428, Sigma) and the triglyceride reagent (TR, T2449, Sigma). Fat bodies from 15-20 wandering L3 larvae were dissected in water, heated to 70°C for 10 min and centrifuged for 2 min at 13000 RPM. Protein concentration was measured by colorimetric method (Pierce™ BCA Protein Assay Kit, Thermo Fisher Scientific). Flat-bottom 96-wells plates containing 20 µL of sample supernatants (in 4 wells) with either 20 µL of PBS (2 wells) or 20 µL of TR (2 wells) were centrifuged for 3 min at maximum speed and incubated for 30 min at 37°C. After incubation, 100 µL of free glycerol kit reagent were added to each sample and incubated for 10 min. Results were normalized using a standard curve generated after e? serial dilutions of glycerol standard solution. Plates were measured on an Infinite 200 Pro microplate reader (Tecan) at 540nm. For data analysis, the value of the “PBS incubated sample” was first subtracted from the value of the “TR incubated sample” to remove the background. Then the amount of triglycerides was estimated using the glycerol standard curve and divided by the protein concentration for each sample.

### Ecdysone feeding experiments

20 early L3 larvae (96–110LJhours AEL) were incubated in 400 μl of crushed banana supplemented with 20 μl of 20-hydroxyecdysone (H5142, Sigma, dissolved in water) at the final concentration of 0.5 mM for 5 hours. Experiments were repeated thrice.

### RT-qPCR

20–25 dissected fat bodies from mid-L3 or wandering L3 larvae were collected in 300 μl of Trizol (Invitrogen). Total RNA was extracted according to the manufacturer’s instructions. RNA quality and quantity were determined using a NanoDrop ND-1000 spectrophotometer. 500 ng of RNA was used to generate cDNA using SuperScript II (Invitrogen, Carlsbad, California, United States). qPCR was performed using dsDNA dye SYBR Green I (Roche Diagnostics, Basel, Switzerland). Expression values were normalized to *rpL32*. Primer sequences used are provided in Table S1.

### Live imaging, staining and microscopy

For Nile red and Bodipy™ staining (to observe intracellular lipid droplets), fat bodies from wandering L3 larvae were dissected in PBS and fixed for 30 min. For Nile Red staining, fixed fat bodies were incubated in Nile Red solution (diluted 1 100 in 1X PBS from a 100 µg/ml stock solution in acetone, Sigma N3013) for 60 min and washed with 1X PBS before mounting. For Bodipy™ staining, fat bodies were incubated for 30 min in Bodipy™ 493/503 (diluted 1 100 in 1X PBS, Life technologies, D3922).

For PH3 staining of hemocytes, mid-L3 larvae were bled in 1X PBS on slides and allowed to adhere for 15 min. Cells were fixed for 20 minutes in PBS, 0.1% Tween 20 (PBT), and 4% paraformaldehyde; then stained with primary antibody [1/500 rabbit anti-PH3 (Upstate/Millipore)]. Secondary staining was performed with Alexa488 anti-rabbit antibodies (Invitrogen). Then cells were stained with a 1/15000 dilution of 4’,6-diamidino-2-phenylindole (DAPI) (Sigma) and mounted in antifading agent Citifluor AF1 (Citifluor Ltd.). The mitotic index was determined by counting the number of PH3 positive cells (green) over all hemocytes (marked with live *hmlΔDsRed.nls*). More than 500 cells were counted per genotype and per assay. The mean and S.E.M. of mitoses per hemocyte pool are shown for each genotype or treatment in the graphs.

For EdU labeling, mid-L3 larvae were fed on fly food containing 1 mM 5-ethynyl-2 deoxyuridine (Click-iT EdU, Thermo Fisher Scientific) for 4 hours for hemocyte staining and 3 hours for lymph gland staining at 25°C. Click-iT EdU staining was performed on released hemocytes or dissected lymph glands according to the manufacturer’s instructions. Counts were performed by visual inspection. More than 500 cells were counted per genotype and per assay for peripheral hemocytes.

For the visualization of NimB5-RFP on hemocytes, larvae were bled without vortexing. Hemocytes were allowed to adhere to slides for 45 min, then were fixed with PFA 4%/Triton0,1% and stained for DAPI (dilution 1/80000, Thermo Fisher Scientific D1306). Images were acquired with a Zeiss AxioImager Z.1, with a 63X objective. Images were taken on a Zeiss LSM 700 Upright confocal microscope at the “Bioimaging and optics platform” in EPFL. Images were processed using Image J.

For the visualization of NimB5-GFP on hemocytes, cells were prepared with the same protocol. Images were taken on a Zeiss LSM 700 Upright confocal microscope at the “BioImaging and Optics Platform” at EPFL. Images were processed using ImageJ.

### Cell area measurement upon spreading

To measure hemocyte area upon spreading, we proceeded as for hemocyte counting. *hmlΔDsRed.nls* hemocytes were let to spread on a glass slide for 45 minutes in Schneider medium supplemented with PTU. After 45 min, cells were fixed with PFA 4%/Triton 0,1% for 15 min, stained with Alexa Fluor™ 488 phalloidin for 3h at a dilution of 1/100 (Thermo Fisher Scientific, A12379) and DAPI at a dilution of 1/80000 for 10min. Cells were imaged under a Zeiss AxioImager Z.1 with a 20X objective and images were analyzed with the Cell profiler software. We design software to select DAPI positive and *hmlΔDsRed.nls* positive cells and to measure their respective GFP area staining. All settings were designed by visual inspection. About 1000 cells per genotype pooled from three independent experiments were analysed. To measure surface area in free floating hemocytes, third instar larvae were bled into PBS w/o calcium and magnesium, supplemented with EDTA 5mM at 37°C for 15 min and placed on ice until analysis. The Invitrogen™ Tali™ Image-based Cytometer was used to measure size of about 10 000 cells.

### *Ex vivo* larval hemocyte phagocytosis assay

*Ex vivo* phagocytosis assay was performed with pHrodo™ Green succinimidyl ester (SE) labeled bacteria (Gram-negative, *Escherichia coli*, and Gram-positive, *Staphylococcus aureus*, Invitrogen) based on the following method [68]. pHrodo™ dye fluorescence increases strongly when submitted to a low pH, so that it allows to follow the internalization process. Five wandering third instar larvae were bled into 120 µL Schneider medium (Gibco) containing 1 nM phenylthiourea (PTU, Sigma). Hemocytes were incubated in 1.5 mL LoBind tubes (Eppendorf) for 45 min at RT at a ratio of 10^5^ pHrodo™ particules/hemocyte. To stop phagocytosis, tubes were kept on ice until FACS analysis.

Phagocytosis of pHrodo™ particles was quantified using a flow cytometer (BD Accuri C6 flow cytometer, USA). 75 µL volume was read in ultra-low attachment 96-well flat bottom plates (Costar no. 3474, Corning) at medium speed (35 µL /min). In a first step, hemocytes were identified using the *hmlΔDsRed.nls* live staining. The fluorescence intensity of single hemocytes was measured in the red channel with 488nm laser and 585/40 standard filter. The green pHrodo™ signal, indicative of hemocytes with effective phagocytosis, was monitored with 488nm laser and 533/30 standard filter. At least 2000 cells per genotype and per assay were analyzed. Results are an average of three independent experiments.

### Crystal cell counts in whole hemocyte population

To determinate the ratio of crystal cells among the total population of hemocytes, we generated wild type and *NimB5^SK1^* fly lines containing the *Lz-Gal4,UAS-GFP* reporter specific for crystal cells with *HmlΔDsRed.nls* that marks both plasmatocytes and crystal cells. For each experiment, eight wandering third instar larvae were bled into 30 µL Schneider medium (Gibco) containing 1 nM PTU and 0.5% PFA to block crystal cell rupture. Larvae were bled individually into 12 wells slides coated with teflon/silane (Tekdon™). Cells were allowed to spread on slides for 30min, then blocked with 4% PFA and 0.1% triton and stained with DAPI. We counted by eye and calculated the ratio of all *Lz-Gal4,UAS-GFP* positive cells over *HmlΔDsRed.nls* positive cells. Results are expressed as an average of 12 wells and are representative of three independent experiments.

### Melanization assays

Phenoloxidase assay: hemolymph was collected by dissecting larvae in 4 °C PBS. The protein concentration was adjusted after a BCA test. Sample volumes were adjusted in 20 μl of 5 mM CaCl_2_ solution. After addition of 80 μl L-DOPA solution (20 mM, pH 6.6), the samples were incubated at 29 °C in the dark. The OD at 492 nm was then measured for 100 minutes using an Infinite 200 Pro microplate reader (Tecan). Reading was performed every 2 minutes after 2 seconds of shaking. An L-DOPA solution without hemolymph was used as a blank. Each experiment was repeated three times. Wild-type and *NimB5^sk1^* third instar larvae were pricked between tracheae on the posterior side with a sterile needle (diameter ~5 μm). Larvae were transferred to 29°C for 30min. Pictures were captured with a Leica DFC300FX camera and Leica Application Suite. For publication purposes, brightness and contrast were increased on some images. For the visualization of sessile crystal cells, twenty third instar larvae were heated in 0.5 ml PBS in Eppendorf tubes for 15 min at 67 °C. Larvae were mounted on glass slides over a white background and imaged. For quantification, black punctae were counted circumferentially in the posterior-most segments A6, A7 and A8.

### Wasp infestation and quantification of fly survival to wasp infestation

For wasp infestations, 25 synchronized second-instar wild-type or *nimB5* mutant larvae were placed on a pea-sized mound of fly food within a custom-built wasp trap in the presence of three female wasps for 2 h (*L. boulardi*). For survival experiments, parasitized larvae were kept at room temperature and scored daily for flies and wasps. The difference between enclosed flies and wasps to the initial number of larvae was set as dead larvae/pupae.

### Statistical tests

Experiments were repeated at least three times on separate days. Unless otherwise indicated, error bars represent the standard error of replicate experiments. Data were analyzed in GraphPad Prism 6.0.

For hemocyte counts, data successfully passed a Shapiro-Wilk normality test (alpha=0.05, n=10) so that we could assume that samples followed a Gaussian distribution. Significance tests were performed using the Student *t* test (with a confidence level of 95%). P values of < 0.05 = *, < 0.01 = ** and < 0.001 = ***. For experiments with more than 2 data sets, significance was tested using Anova (with a confidence level of 95%). P values of < 0.05 = *, < 0.01 = ** and < 0.001 = ***

## Authors’ contribution

Conceived and designed the experiments: ER BL. Performed the experiments: ER BP JPD JPB MP SK. Analyzed the data: ER BP JPD BL. Wrote the paper: ER BL.

## Acknowledgements

We thank the VDCR in Vienna and the Bloomington Stock Center (NIH P40OD018537) at Indiana University for the fly stocks. We thank the BIOP platform (EPFL) for help with using the confocal microscope and design of Cell profiler analysis. We thank the BioEM platform (EPFL) for help with electron microscopy experiments. We thank Marie Meister, Claudine Neyen and Mark Hanson for critical reading of the manuscript.

## Supplementary figure legends

**Figure S1. Flies with high number of hemocytes have a normal “inflammatory state”**

**(A - C)** RT-qPCR quantification of *diptericin* (**A**), *drosomycin* (**B**) and *upd3* (**C**) genes in fat body extracted from *hml^Δ^* > *w* (WT), *hml^Δ^*> *pvf2, hml^Δ^* > *ras* IR and *hml^Δ^* > *ras^v12^* flies, in unchallenged conditions (**A, B** and **C**) and 3 hours post-infection with ECC15 (**A**) or *M.luteus* (**B**). **(D, E)** Phagocytic index of hemocytes extracted from *hml^Δ^* > *w* (WT), *hml^Δ^*> *pvf2, hml^Δ^* > *ras* IR and *hml^Δ^* > *ras^v12^* flies, using green pHrodo particles coupled with *S. aureus* (**D**) and *E. coli* (**E**). *eater* mutant is defective for Gram positive bacteria internalization and is used as a negative control.

**Figure S2. Hemocytes number is affected by nutrients intake**

**(A)** Total counts of peripheral hemocytes from *w^1118^* and *crq_Δ_* mutant at L3 wandering larval stage. Animals are raised on normal diet, **(B)** Larvae with identical genotypes were dissected and fat body was Bodipy stained. Scale bars = 58 μm hemocytes from *Hml^Δ^-GAL4,UAS-GFP* L3 wandering larvae fed on normal diet (ND), poor diet (PD), normal diet supplemented with glucose (ND+glucose) and normal diet supplemented with lard (HFD-lard 6%). (**D**) RT-qPCR quantification of *NimB5* over *Rpl32* transcripts from fat bodies of mid-L3 larvae raised on agar medium and dissected at indicated time points. *P* values from Anova test for **C** and **D**.

**Figure S3. NimB5 is secreted into the hemolymph.**

**(A)** Western blot analysis with hemolymph extract from L3 wandering larvae over-expressing *UAS-NimB5-RFP* with the driver *lpp^TS^-Gal4* reveals the presence of the RFP fusion protein at the expected size of ≈60-65 kDa molecular weight (NimB5: 35.34 kDa and RFP: 27 kDa). **(B)** Image of L3 wandering larvae expressing *UAS-NimB5-RFP* with the *lpp^TS^-Gal4* driver. Tissues (notably fat body) were marked with Tubulin-GFP while NimB5-RFP can be seen in red. NimB5-RFP accumulates in nephrocytes indicating that the protein is secreted into the hemolymph.

**Figure S4. Generation of N*imB5* deficient flies by CRISPR-Cas9 system.**

**(A)** Representation of the *Nimrod* gene locus on the 2^nd^ chromosome (34E5). The *NimB5* gene is mutated in the first exon via a deletion of 13 bp. **(B)** The mutation of *NimB5* gene leads to a premature translation stop resulting in a truncated protein lacking 297 amino acids.

**Figure S5. NimB5 regulates hemocyte proliferation**

**(A)** Peripheral hemocyte counts of *w^1118^* or *NimB5^SK1^* isogenized lines at L3 wandering stage. **(B)** Peripheral hemocyte counts from *wt*, *Df(2L)ED793 / +* and *Df(2L)ED793 / NimB5^SK1^* L3 wandering larvae with the *hml^Δ^.dsred.nls* marker. **(C)** Peripheral hemocyte counts of *C564-GAL4*/*+*; *+/+*, *C564-GAL4/UAS-NimB5-IR*; +/+ and *C564-GAL4/+; UAS-NimB5/+* L3 larvae. **(D)** Peripheral hemocyte counts of *hmlΔGal4,UAS-GFP*/+ (WT), *hmlΔ-Gal4,UAS-GFP*/*UAS-NimB5-IR*; +/+ and *hmlΔGal4,UAS-GFP*/+; *UAS-NimB5*/+ L3 larvae. **(E, F)** Complementation of the *NimB5^SK1^* mutation by over-expressing *NimB5* from either the fat body **(E)** or hemocytes **(F)**. Genotypes are indicated on graphs. All results represent a total of five animals. *P* values from Student’s *t-*test for **A**, **C** or from Anova for **B**, **E** and **F.**

**Figure S6. NimB5 is not required for melanization and encapsulation.**

**(A)** Phagocytic assay using green pHrodo particles. Hemocytes dissected from *WT* or *NimB5^SK1^* L3 wandering larvae show identical phagocytic capacity towards *E. coli* and *S. aureus* conjugated bioparticles. Data were acquired using flow cytometry with gating against red fluorescent cells, then green positive cells. **(B) A** DOPA enzymatic assay reveals that hemolymph from *NimB5^SK1^* L3 wandering larvae has higher melanization activity compared to wild-type larvae. This higher level can be explained by the higher absolute number of hemocytes (and crystal cells) in *NimB5^SK1^* larvae. **(C)** Pricking assay shows identical melanization reaction as monitored by the size of the black spot at the injury site in wild-type and *NimB5^SK1^* larvae. (**D, E**) Representative images of lamellocytes spread on a slide **(D)** or circulating in whole animals **(E).** Larvae carry the *PPO3gal4,uasGFP* insertion, that is specifically expressed in lamellocytes [69]. Hemocytes and larvae were collected two days after *L. boulardi* wasp infestation. Wild-type and *NimB5^SK1^* larvae do not show any difference in their ability to differentiate lamellocytes upon wasp infestation. **(F)** Ratio of *lz-Gal4,UAS-GFP* positive cells (crystal cells) over *hml^Δ^.dsred.nls* positive cells (plasmatocytes). *NimB5^SK1^* L3 wandering larvae show a significantly lower ratio compared to wild-type. **(G, H)** Heating assay reveals a lower number of visible black punctae (crystal cells) below the carcass of *NimB5^SK1^* larvae compared to wild-type larvae. The decreased number of black spots is associated with a decreased number of sessile crystal cells. The decreased number of sessile crystal cells in *NimB5^SK1^* larvae could be a secondary consequence of an adhesion defect of *NimB5^SK1^* hemocytes, as previously described for *Eater* deficient mutants [30]. **(H)** Representative images of data presented in **(G)**. *PPO1, PPO2, PPO3* deficient larvae lacking cellular and hemolymphatic phenoloxidase were used as a control for **B** and **H** [69]. For **F**, *P* values are indicated from Student’s *t-*test. For **G**, *P* values are indicated from Anova test.

**Figure S7. NimB5 mutation does not affect hemocyte shape.**

(A) Width hemocytes measurement from wild-type and *NimB5^SK1^* mutant L3 wandering larvae. Analysis are processed with Tali® Image-Based Cytometer.

**Figure S8. NimB5 is required to sustain wasp development during infestation.**

*Leptopilina boulardi* infestation in *wild-type* and *NimB5* flies with a yellow-white genetic background (**, *P* >O.0071; chi-square = 9,885; df = 2). Results are represented as a sum of 120 animals for each genotype in six different experiments. In this experiment, we monitored the three possible outcomes of infestation: 1) wasp-infested individuals die as larvae or as pupae (both wasp and fly die), 2) a wasp emerges from the pupa (wasp success), or 3) a fly emerges from the infested pupa (fly success). This experiment shows that the endoparasitoid wasp *L. boulardi* is less successful to infect *NimB5* deficient mutants compared to wild-type. This is likely due to increased hemocyte number of *NimB5* larvae that either improves the cellular response (i.e. encapsulation) against the parasitoids or depletes lipid store. A depletion of lipid store would explain why we observe increased lethality at the pupal stage corresponding to case 1 (both wasp and fly die).

**Figure S9. NimB5 may interact with other(s) protein(s) than Eater.**

Peripheral (sessile and circulating) hemocyte counts of *w^1118^*, *NimB5^SK1^*, *eater^1^* and *NimB5^SK1^*; *eater^1^* animals combined with *hml^Δ^*.*dsred.nls* from L3 wandering larvae. *P* values are calculated from Anova test.

**Figure S10. Model of NimB5 role in the control of hemocyte function.**

NimB5 expression adjusts the growth of the peripheral hematopoietic compartment to the metabolic state of the host. NimB5 is produced by the fat body upon starvation or during metamorphosis, and subsequently secreted into the hemolymph. NimB5 binds to hemocytes to promote adhesion to the cuticle (black arrows) and possibly contact between cells (red arrowheads).

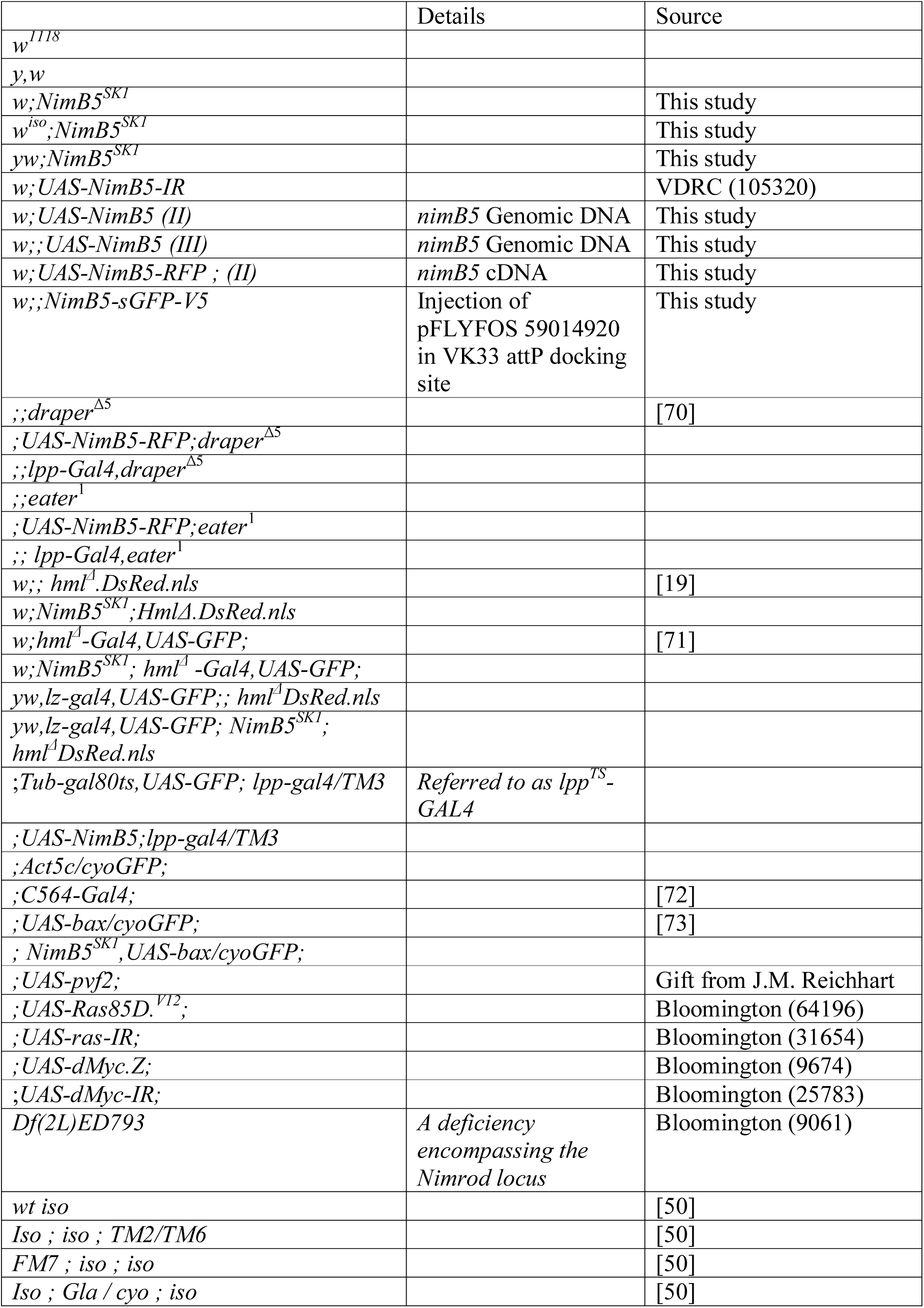
List of stocks generated and used in this manuscript.

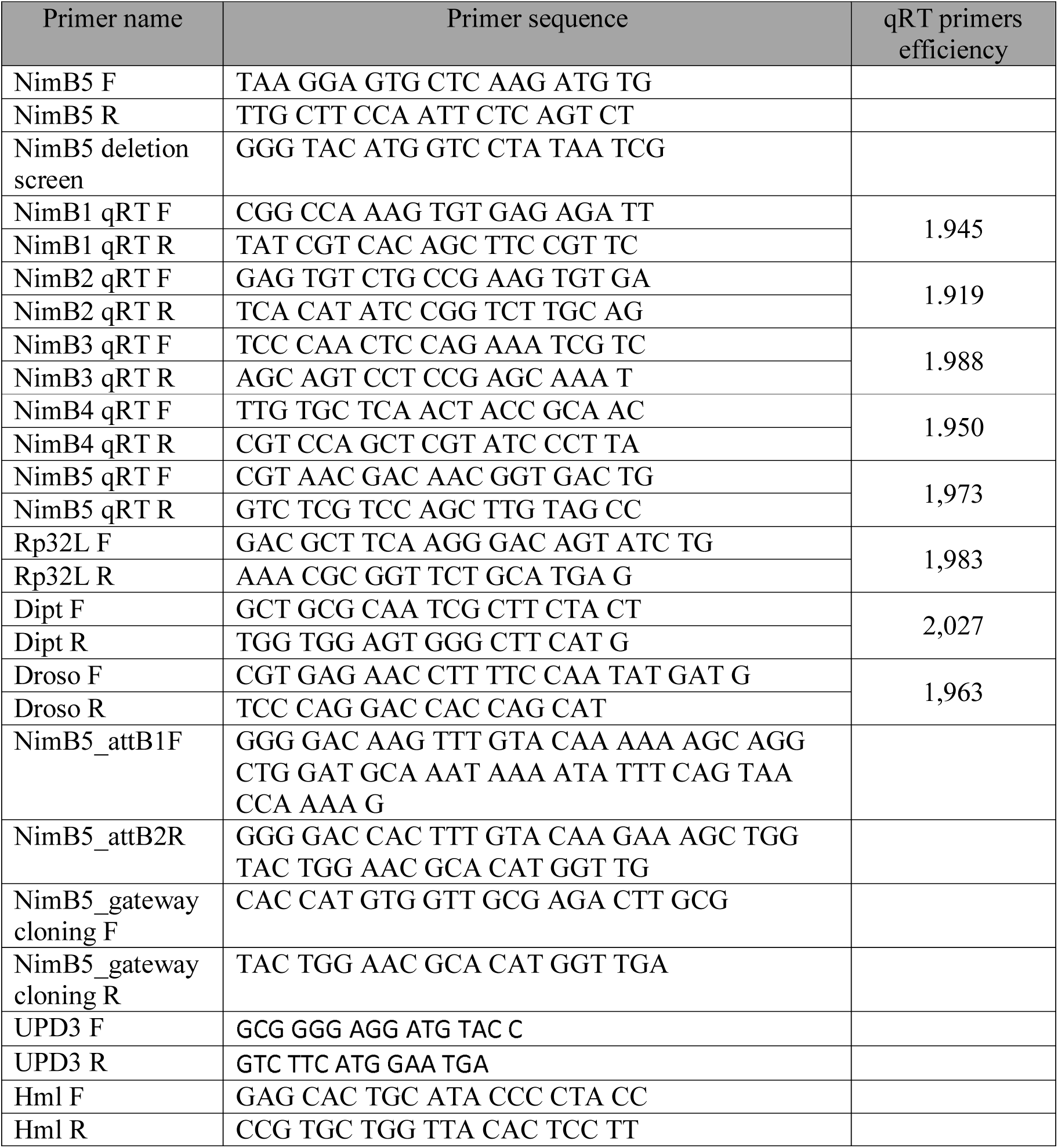
List of primers used in this manuscript.

List of plasmids and BAC used in this manuscript.

List of plasmids and BAC used in this manuscript
| Plasmid name | Antibiotics selection | Origin |
| --- | --- | --- |
| pENTR/D-TOPO | Kan<sup>R</sup> | Invitrogen |
| pTW | Amp<sup>R</sup> | DGRC |
| pTWR | Amp<sup>R</sup> | DGRC |
| pOT2-CG16873<br>Full lenght cDNA clone | Cm<sup>R</sup> | DGRC (IP09831) |
| pCaSpeR4 | Amp<sup>R</sup> | Our collection |
| BACR09N24 (RP98-9N24) | Cm<sup>R</sup> | BACPAC Resources Center |

