## Supplementary figures and images for "Metabolic adjustment of *Drosophila* hemocyte number and sessility by an adipokine"

### Supplementary figure S1

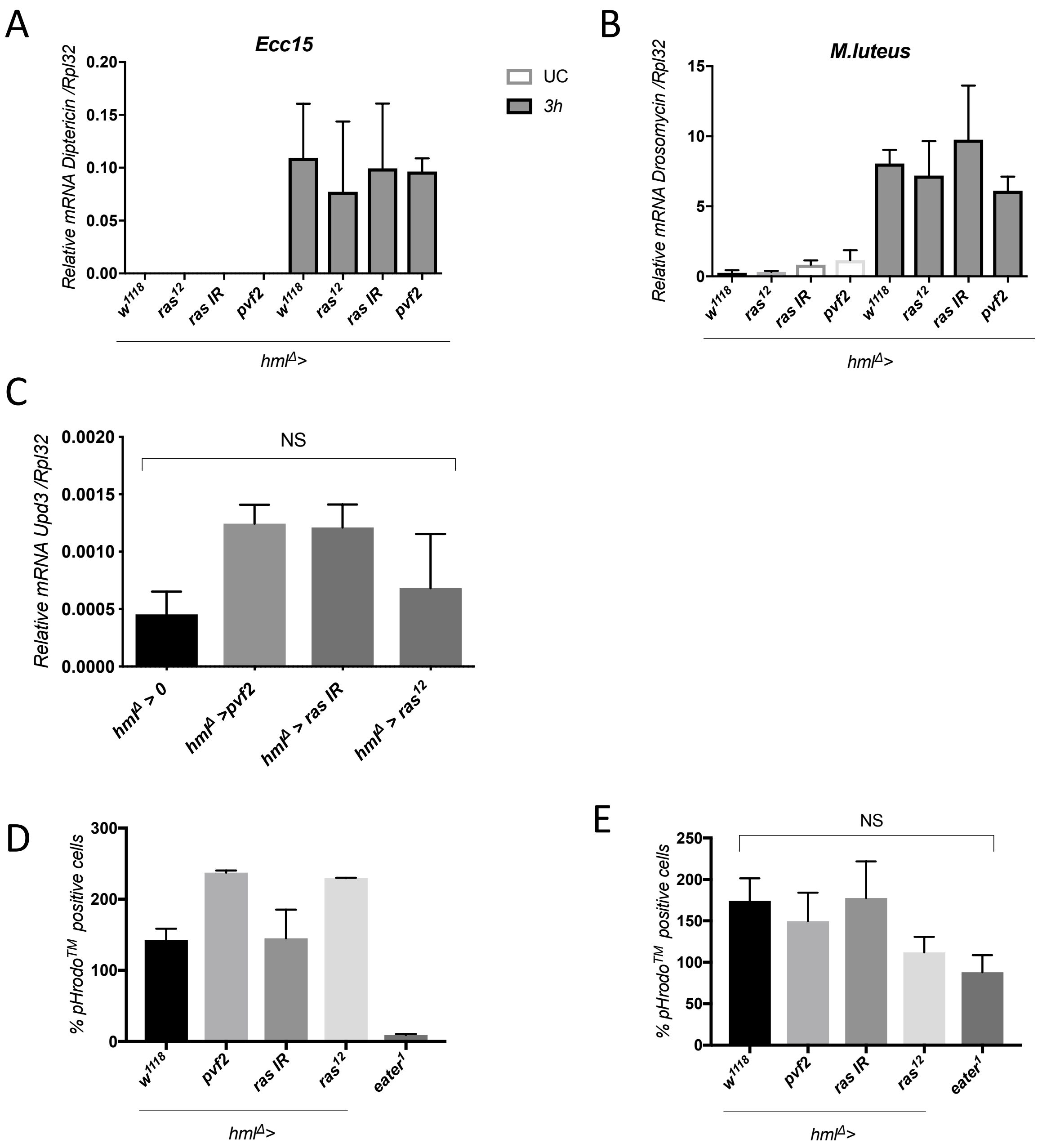


Figure S1 Ramond et al.

### Supplementary figure S2

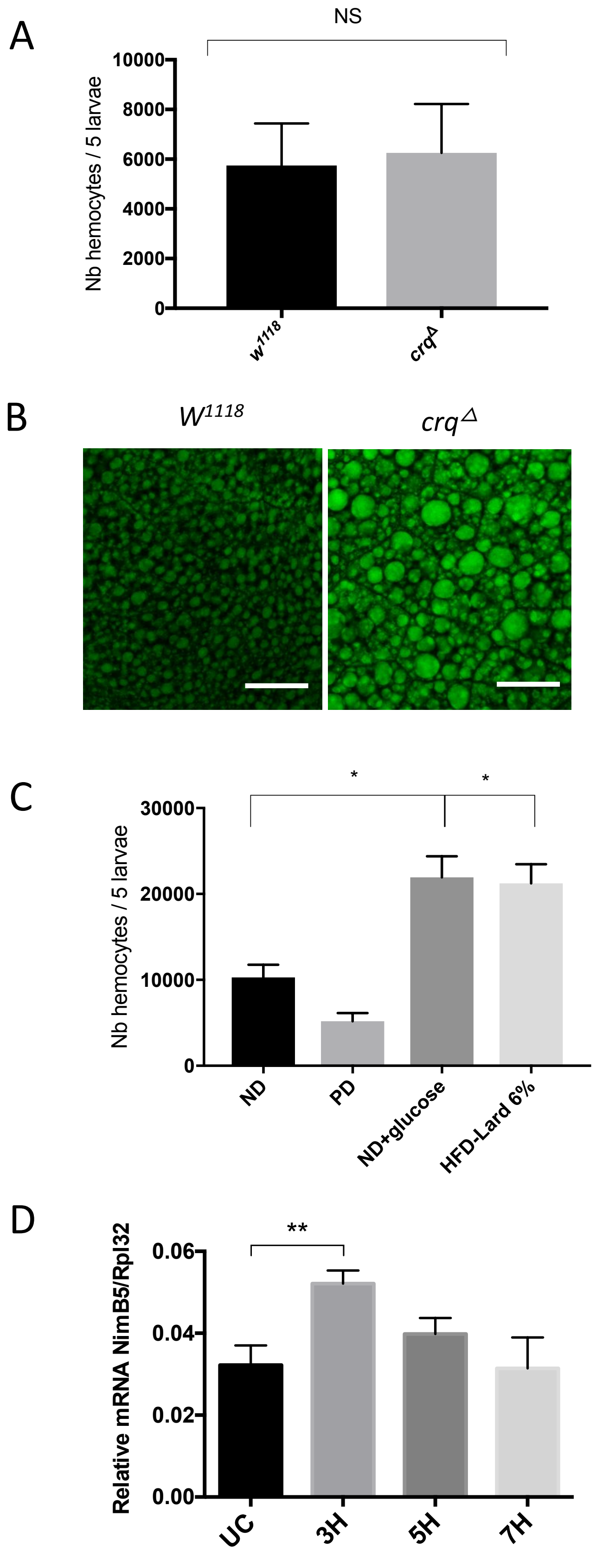


Figure S2 Ramond et al.

### Supplementary figure S3

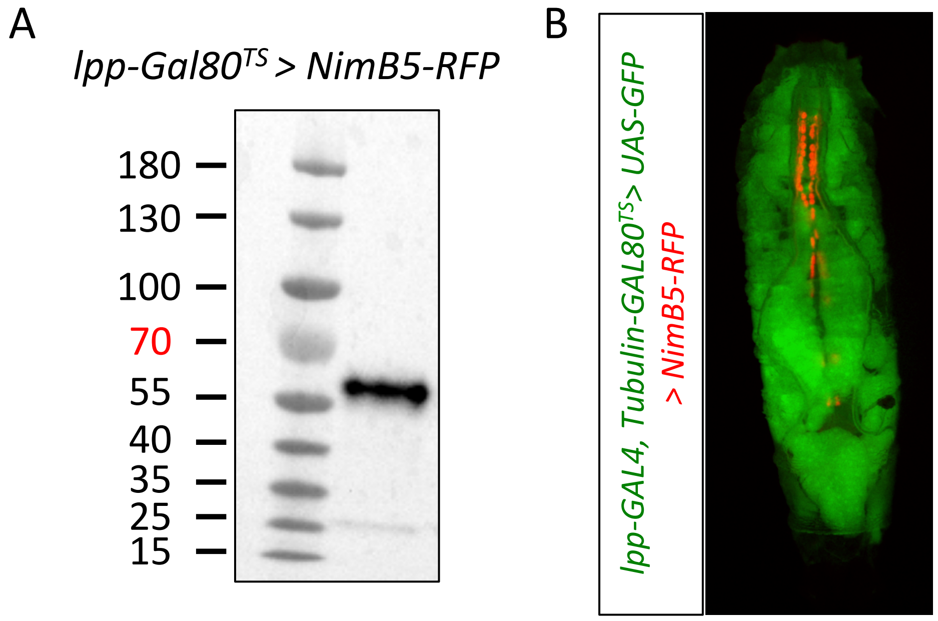


Figure S3 Ramond et al.

### Supplementary figure S5

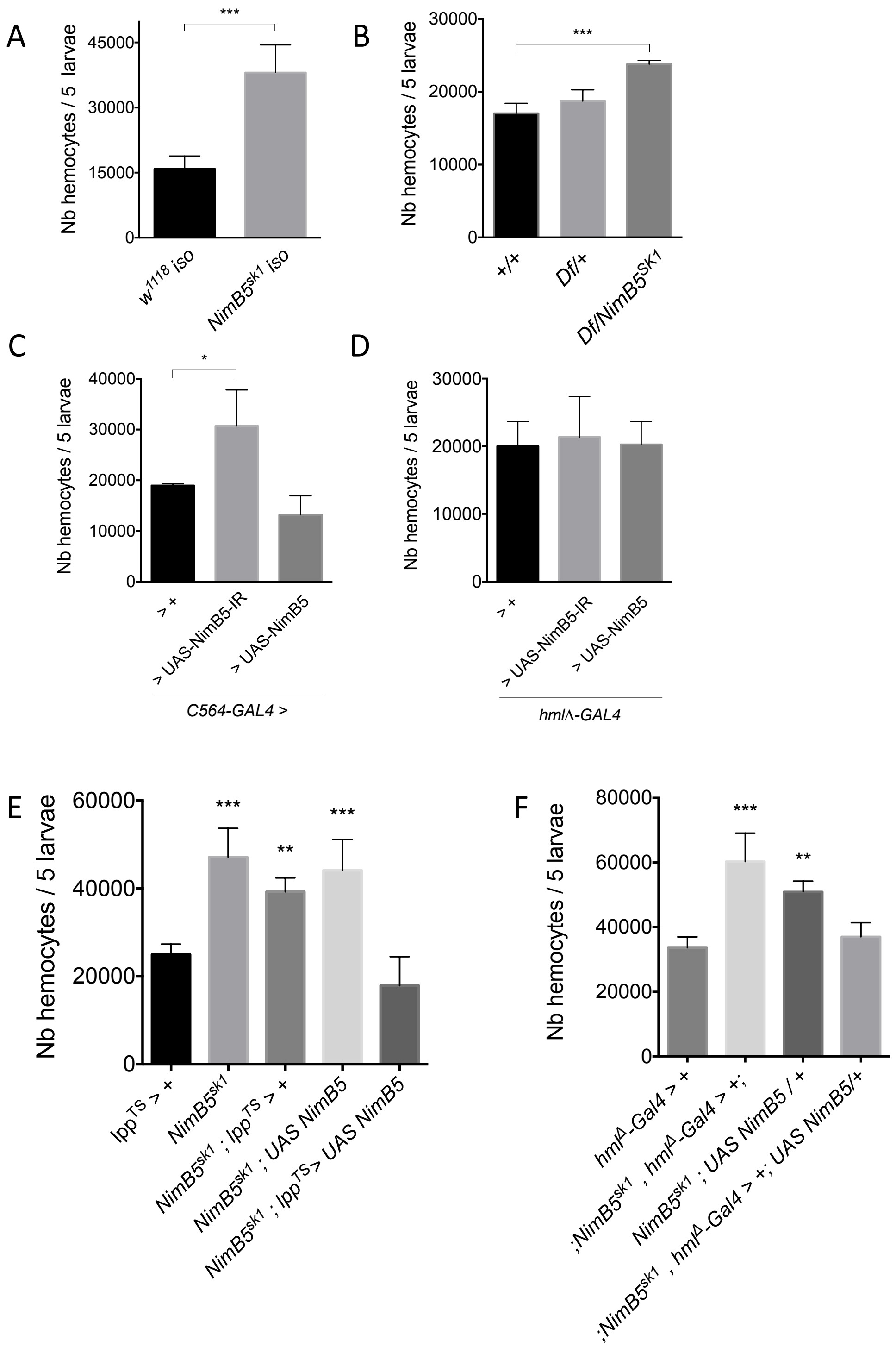


Fig S5 Ramond et al.

### Supplementary figure S6

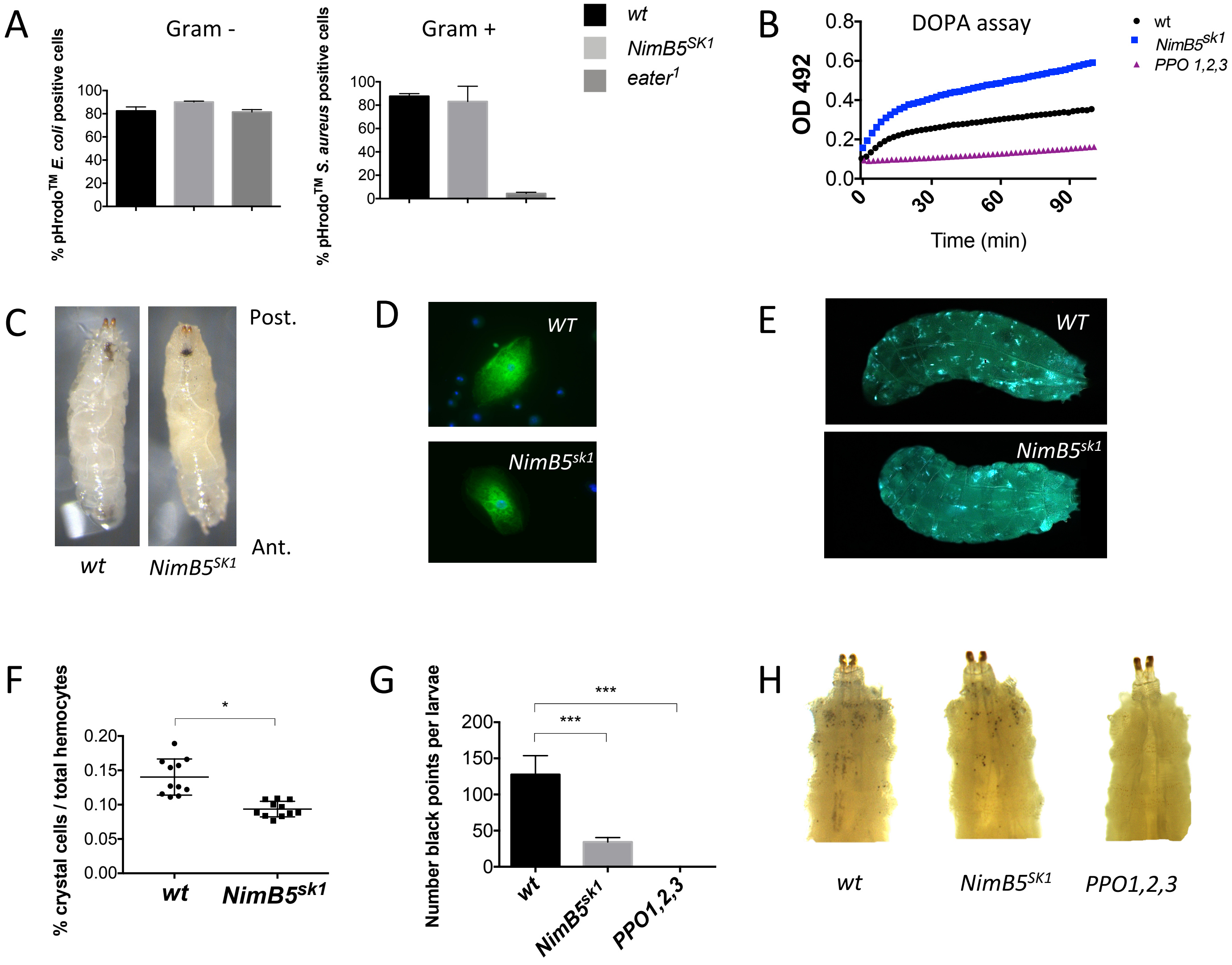


Fig S6 Ramond et al.

### Supplementary figure S7

A

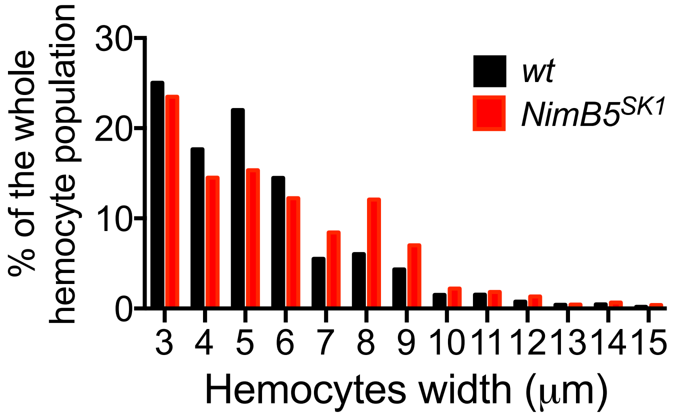


Fig S7 Ramond et al.

### Supplementary figure S8

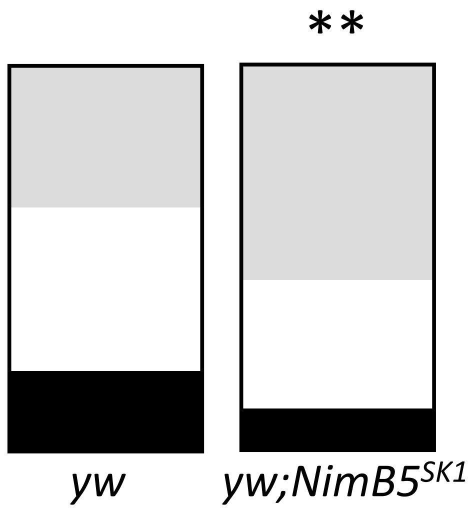


Fig S8 Ramond et al.

### Supplementary figure S9

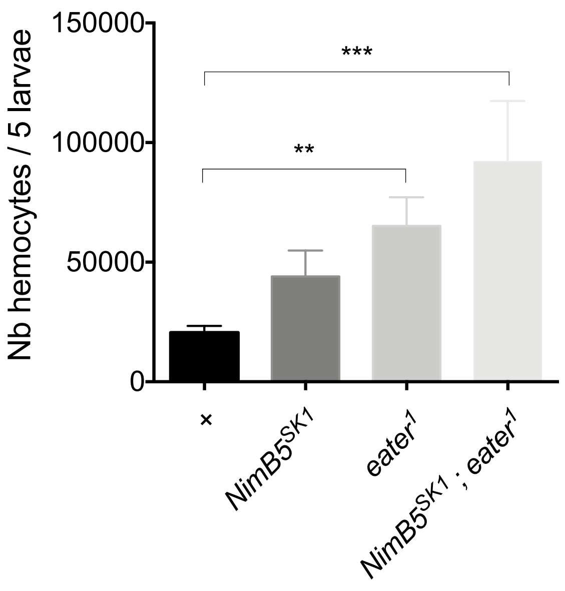


Fig S9 Ramond et al.

### Supplementary figure S10

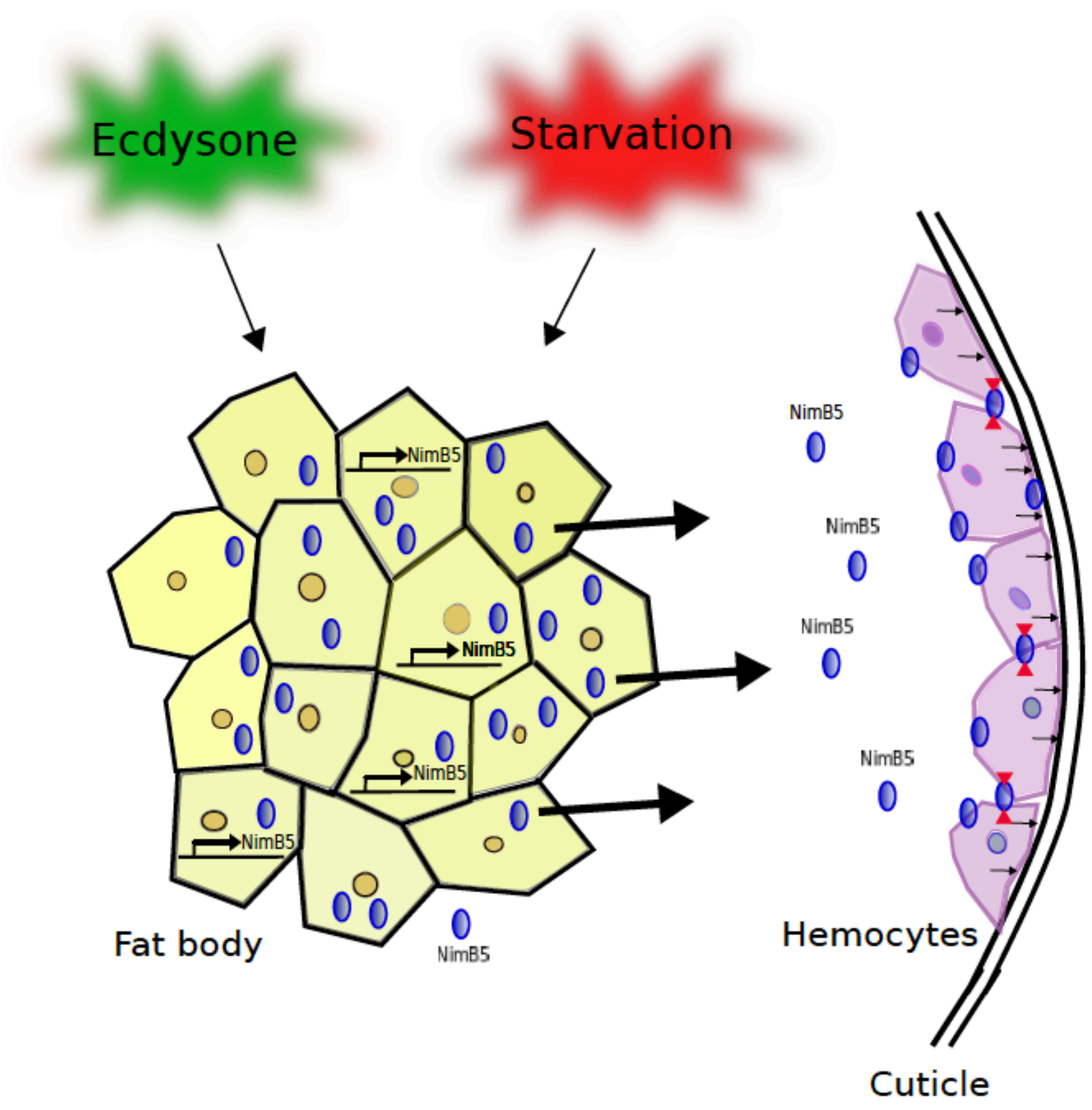


Fig S10 Ramond et al.
