## Supplementary figure S4 for "Metabolic adjustment of *Drosophila* hemocyte number and sessility by an adipokine"

A

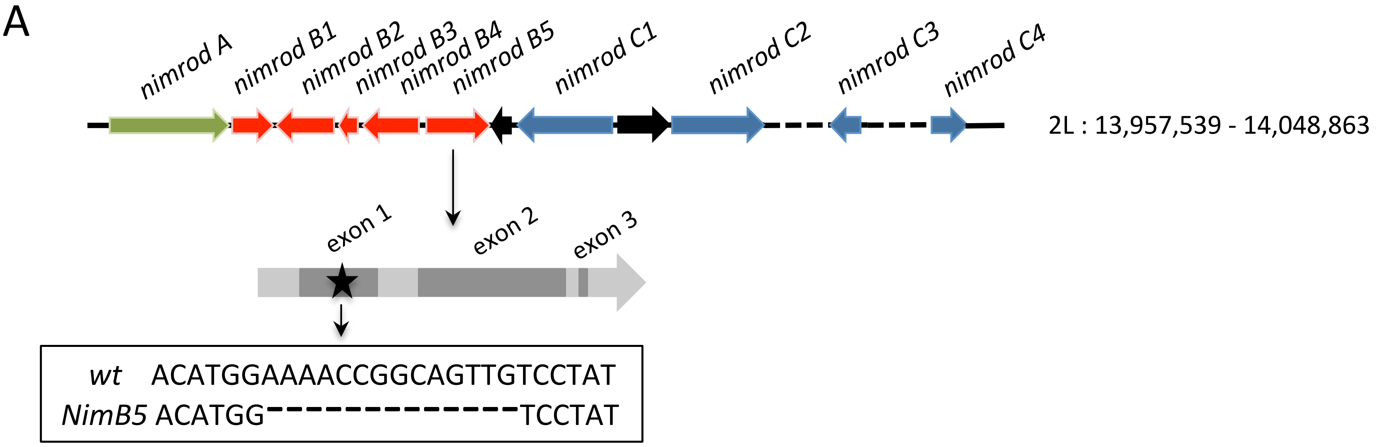


B

### Sequence of the NimB5 protein

**M** W L R D L R I R L S F F H L M V C L P S I Y T N V P S Y S D R
Y N R R P V S Q D P G M G T Y G R G Y M E N R Q L S Y N R P
G P A N F Q D P R N V E L Q P K R R E Y L I R S H E T Q S D R
G Q H K C R I W V P P D T V E K Y S Y P S V I Q T D Q A N R L
S L I E V C C T G Y S A S R L M G V T V C R A Q C G C Q N G S
C K I P G E C E C Y D G F V R N D N G D C V F A C P L G C Q N
G Q C Y L D G S C Q C D P G Y K L D E T R R F C R P I C S S G C
G S S P R H N C T E P E I C G C S K G Y Q L T D D G C Q P V C
E P D C G I G G L C K D N N Q C D C A P G Y N L R D G V C Q
A D C Y Q K C N N G V C V S R N R C L C D P G Y T Y H E Q S T
M C V P V **Stop**

### Sequence of the truncated NimB5$^{SK1}$ protein

**M** W L R D L R I R L S F F H L M V C L P S I Y T N V P S Y S D R
Y N R R P V S Q D P G M G T Y G R G Y M V L **Stop**

Fig S4 Ramond et al.
